# HyperPRI: A Dataset of Hyperspectral Images for Underground Plant Root Study

**DOI:** 10.1101/2023.09.29.559614

**Authors:** Spencer J. Chang, Ritesh Chowdhry, Yangyang Song, Tomas Mejia, Anna Hampton, Shelby Kucharski, TM Sazzad, Yuxuan Zhang, Sanjeev J. Koppal, Chris H. Wilson, Stefan Gerber, Barry Tillman, Marcio F. R. Resende, William M. Hammond, Alina Zare

## Abstract

Collecting and analyzing hyperspectral imagery (HSI) of plant roots over time can enhance our understanding of their function, responses to environmental factors, turnover, and relationship with the rhizosphere. Current belowground red-green-blue (RGB) root imaging studies infer such functions from physical properties like root length, volume, and surface area. HSI provides a more complete spectral perspective of plants by capturing a high-resolution spectral signature of plant parts, which have extended studies beyond physical properties to include physiological properties, chemical composition, and phytopathology. Understanding crop plants’ physical, physiological, and chemical properties enables researchers to determine high-yielding, drought-resilient genotypes that can withstand climate changes and sustain future population needs. However, most HSI plant studies use cameras positioned above ground, and thus, similar belowground advances are urgently needed. One reason for the sparsity of belowground HSI studies is that root features often have limited distinguishing reflectance intensities compared to surrounding soil, potentially rendering conventional image analysis methods ineffective. Here we present HyperPRI, a novel dataset containing RGB and HSI data for in situ, non-destructive, underground plant root analysis using ML tools. HyperPRI contains images of plant roots grown in rhizoboxes for two annual crop species – peanut (*Arachis hypogaea*) and sweet corn (*Zea mays*). Drought conditions are simulated once, and the boxes are imaged and weighed on select days across two months. Along with the images, we provide hand-labeled semantic masks and imaging environment metadata. Additionally, we present baselines for root segmentation on this dataset and draw comparisons between methods that focus on spatial, spectral, and spatial-spectral features to predict the pixel-wise labels. Results demonstrate that combining HyperPRI’s hyperspectral and spatial information improves semantic segmentation of target objects.

## 1. Introduction

Researchers have studied the links between root traits and root function for the purposes of phenotyping and guiding plant choices in agriculture [1, 2, 3]. A high-priority task is identifying genotypes and characteristics of high-yield, stress-tolerant crops that can sustain the projected nutritional needs of 10 billion people by 2050 [4]. Studies have ranged across multiple plants such as cassava [5], maize [6], chicory [7], soybean [8, 9], cotton [10], wheat [11], and many other plants [12], and the number of studies continues to grow each year [3]. Phenotypic traits such as root length, volume, surface area, and count are utilized as features to understand the roots’ functions. Studies have also related certain physical root characteristics to genotypes that have better drought resilience [5, 13]. The use of RGB data to do high-throughput plant phenotyping with rhizoboxes has led to the creation of rhizobox environments and computer hardware and software that enable quicker and better phenotyping experimentation and evaluation [14, 15, 16, 9, 17, 10, 18]. However, the above RGB studies provide an incomplete view of a plant’s root physiology and root function, and while there has been recent success in predicting belowground root traits using aboveground HSI data for peanuts [19], it is not always clear that a given plant’s aboveground status during drought is indicative of its belowground status [20]. This calls for more analysis of belowground traits to gain a more complete picture of a plant’s drought response.

Current methods of acquiring root traits are both destructive and non-destructive. Destructive methods include digging up plants from their original placement to image in 2D or 3D settings [21, 13]. A common non-destructive method is the use of minirhizotrons (MRs) [1, 2, 12]. To be minimally invasive, MR tubes are inserted into the soil before a plant grows its roots so that root traits may be collected over time. An alternative to MRs is the rhizotron or rhizobox method that involves growing plants within boxes made of clear material so that root traits may be monitored over time [14, 16, 22, 17]. Other non-destructive methods include tomography scans, X-ray CT scans [23], and MRIs to see beneath the soil surface [24]. As a final note, many non-destructive methods depend on the use of low-cost, low-resolution hardware that is deployed at scale. Such a constraint gives rise to the need for techniques of enhancing image resolution so that root trait analysis may be improved [25, 26].

While tomography, X-ray CTs, and MRIs can reveal more types of roots belowground, they also require a tremendous amount of resources, preventing their deployment in high-throughput applications. X-ray CTs also have the risk of altering or impeding plant growth [27, 28]. Besides them, each of the other references above uses RGB imaging to infer root physiology. The referenced rhizobox imaging studies tend to focus on phenotyping through an analysis of the root system architecture, but the physical characteristics of root length, volume, surface area, and count lack a complete view of a plant’s status and root function. Studying hyperspectral, phenotypic properties invisible to the naked eye can provide more information on plant physiology (ie. chemical traits [29]). Recently, in relation to phytopathology, researchers were able to use plant leaf spectra to classify healthy and blighted potatoes [30], while others have used HSI data to study a fungi infecting southwestern white pines [31]. For roots specifically, studies have utilized near-infrared (NIR) images and HSI [32, 33] to improve root component classification and root phenotyping. Nonetheless, we were unable to find a publicly available visible-NIR or HSI plant root datasets that would allow further investigation into the relationship between root spectra and its surrounding rhizosphere.

In this work, we present a HSI rhizobox dataset containing both RGB and HSI data along with fully annotated masks for selecting root and soil pixels. Additionally, we monitor 64 boxes for up to 15 timesteps across two months, and most of the plants go through a natural dehydration and rewatering process (ie. some were in a control group). The dataset’s hyperspectral (HS) images over time can reveal a deeper understanding of the relationship between root characteristics and root function. By simulating drought conditions, we have acquired numerous spectral signatures that span the wettest to near-driest soil conditions. In our experience, the two extremes for soil condition also represent the easiest (ie. wettest) and most difficult segmentation environments (ie. near-driest). We provide images for studying root traits in both peanut and sweet corn roots, and we apply a subset of peanut images to the task of semantic segmentation for both types of image data. Our dataset contains more in-depth evaluation of plant roots across a broad range of soil conditions that can be applied to studies on phenotyping prediction [33, 34], dense root systems [35, 36], interactive dynamics between root and rhizosphere [37], and drought resiliency [5, 6]. Most recently, HyperPRI has been used for studying and comparing the crop plants’ drought tolerance and recovery [38]. The data can also be applied to the tasks of data reconstruction and semantic segmentation. In this work, we demonstrate that root segmentation is improved with the addition of root and soil spectral signatures.

## 2. Related Datasets

Current methods of acquiring root traits and carrying out root phenotyping are either destructive or non-destructive [13]. MR technology is a non-destructive method that has been used for multiple decades to do insitu monitoring of plant roots [1, 2]. Existing, public MR datasets include RGB root images for soybean [8], chicory [7], and multiple other plants [12]. Researchers have also measured root traits and conducted root phenotyping through rhizotron/box laboratory setups [14, 16, 22, 39], but the imaged rhizoboxes are not widely available online. Compared to MR, rhizoboxes give more root information by providing a full view of the root system architecture. In addition to the above, there is a database containing multiple root system datasets [40, 41], but many either are not pictured in-situ or do not have public ground-truth annotation masks of the root systems.

Although RGB MR or rhizobox studies are becoming more common [14, 16, 22, 17, 39], there are few methods that add spectral reflectance (HSI data) to the list of root traits [42]. In particular, a prior rhizobox study used spectral information to better differentiate between root classes (ie. dead, living, rotting, etc) [32]. Two more recent studies used spectral signatures to improve root segmentation and to demonstrate that invisible wavelengths of HSI data can contribute to better root phenotyping [33, 34]. However, neither of these two studies have made their data publicly available, and the former of the two studies captures wavelengths between 900 and 1700 nm [33] and covers a different wavelength range and larger spectral width (3.1 nm) than our dataset. There are also no datasets that provide natural dehydration and rewatering processes, and we did not find any RGB or HSI datasets that provide multiple rhizosphere image timesteps for monitoring roots from seedling to reproductively mature plants.

In HyperPRI, we provide RGB and HSI rhizobox images acquired across multiple days over the course of two months. Our dataset includes fully annotated segmentation masks for all root images and box weight metadata per each imaged day. The metadata is a series of box weights (in grams) that includes the empty box, the box with dry soil, the box with wet soil, and the box weight at each monitored day. Box weights for each monitored day are limited to those that remained alive and intact for the duration of the two months. The dataset will be made available online at doi: 10.1101/2023.09.29.559614v1. Based on our review, there is no other public dataset containing time-series HSI rhizobox images with fully annotated segmentation masks.

## 3. Data Acquisition and Annotation

This section describes the environment and methodology used for acquiring the imaging data and associated metadata in the context of machine learning research. The imaging setup involved using the HinaLea Model 4200 Hyperspectral Imager camera (2018) that images between the wavelengths of 400 and 1000 nm with a spectral width of 2 nm (299 spectral bands) and with a resolution of 968 *×* 608. The camera allowed us to capture both HSI and RGB images of the plant boxes at regular intervals throughout the monitoring period. The plants were closely monitored over a total of two months between June and August.

Our research involved the use of 64 rhizoboxes that measured a height-width-depth of 34.5 *×* 21 *×* 3.8 cm^3^ and covered a region that is 9.3 *×* 14.8 cm^2^ near the soil surface of the rhizobox (each pixel has 0.15 mm spatial width). We constructed the boxes with a clear plastic sheet (Lexan™) in front and filled them with approximately two kilograms of Turface soil, accommodating two species: sweet corn and peanut. The specific growing soil was a fritted calcined clay, Profile Porous Ceramic (Greens Grade™, Turface Athletics, Buffalo Grove, IL, USA). To ensure a consistent distance from the lens while acquiring RGB and HSI data, we used the setup in Figure 1b, which uses a wood block to align the rhizobox into the same position in front of the camera. More information about the genotype and specific box number is in Table 1. Each box was named according to the date of imaging and box number. The naming scheme followed the format yyyymmdd boxNN, where mm denotes the month, dd denotes the day, yyyy denotes the year, and NN denotes the box number. Fig. 1 shows the imaging setup on the right and a sweet corn rhizobox example on the left. The authors primarily annotated the training masks using the RGB images.

**Figure 1:**
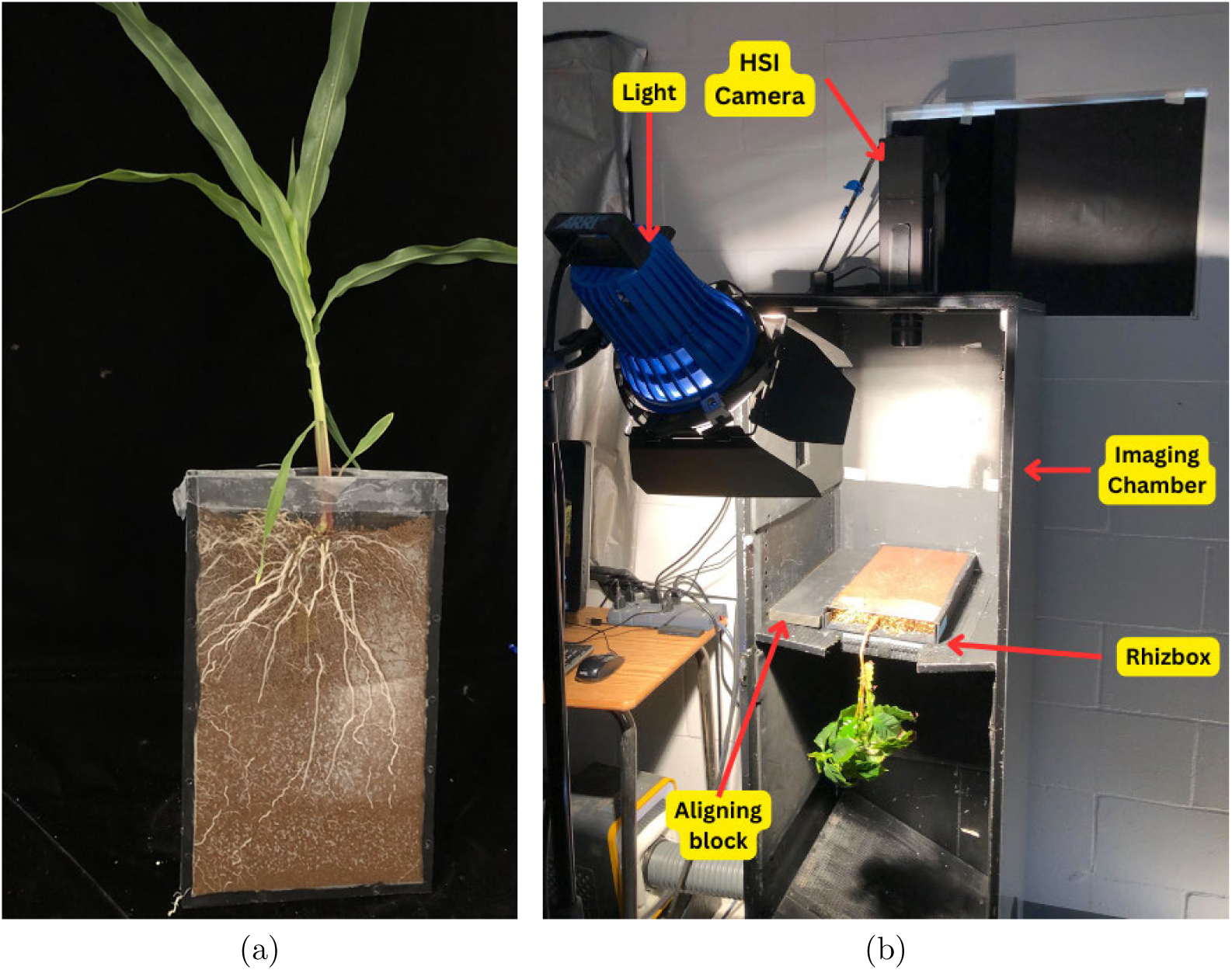
Imaging Setup for HyperPRI Data Acquisition. (a) A sweet corn rhizobox from our data collection on (June 15, 2022), 27 days after planting. (b) Hyperspectral Camera setup that was used to collect this dataset. The HSI camera at the top captures images in the range 400 – 1000 nm, the illuminator light (white light) assists in the capturing of all bands. An aligning block is used to ensure that the rhizobox is always at the same position and the same region is captured.

**Table 1:** Species and genotypes of all the boxes in the dataset.

| Box Numbers | Species | Genotype |
| --- | --- | --- |
| 01 – 16 | Sweet Corn | IL395a |
| 17 – 32 | Sweet Corn | IL14H |
| 33 – 48 | Peanut | TUFRun511 [45] |
| 49 – 64 | Peanut | 10X34-4-4-1-2 |

Fig. 2 shows the timeline that was followed for data collection. The peanut seeds were planted in rhizoboxes on May 26, 2022, and the sweet corn was planted on May 19, 2022 (between Fig. 2b and 2c). A subset of boxes from each species was subjected to a dry down period (16 peanut, 20 sweet corn), while the remaining boxes were regularly watered to serve as controls. For sweet corn, the dry down period occurred from June 28, 2022, to July 21, 2022. The dry down period for peanut took place from August 8, 2022, to August 19, 2022 (between Fig. 2c and 2d). More specifically, sweet corn plant roots were imaged between 26 and 70 days after planting (DAP), and peanut plant roots were imaged between 22 and 83 DAP. Drought was initiated around sweet corn’s V7 – V9 stage at 39 DAP and around peanut’s R6 stage at 75 DAP. At the end of the dry down period, plants are rewatered (Fig. 2e). The annotation methodology involved manual labeling of the images using various tools such as VIA [43, 44], Photoshop, and GIMP. Initially, the annotations were performed by hand, and as the model improved, active annotation techniques were employed to enhance the accuracy and efficiency of the annotation process. The masks have four labels in peanut: 0 - Soil, 1 - Nodules, 2 - Peanut Pegs, 3 - Roots and two in Sweet Corn: 0 - Soil, 3 - Roots. During our data labeling, we found the nodule and peg classes to have little representation and chose to collapse the classes into the simpler task of segmenting roots out of soil. Therefore, for the segmentation purposes, we put the labels 1, 2 and 3 under roots.

**Figure 2:**
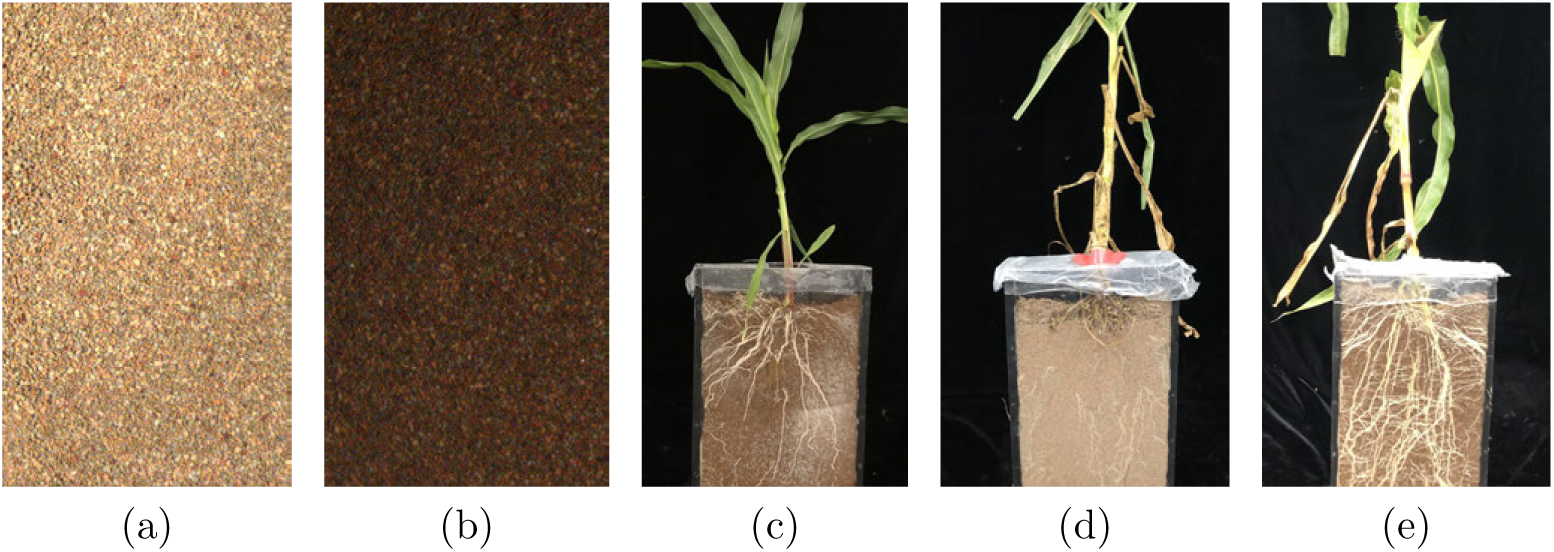
Plant growth time lapse. Shown is a series of five images labeled (a) through (e), capturing pivotal stages in the research project. (a) and (b), obtained using the HSI camera, focus on a specific region within the box. (a) presents dry soil conditions, while (b) captures wet soil. As the roots reached the imaged area (c), regular intervals were established for capturing images to monitor plant growth. Subsequently, a drought initiation phase was implemented, wherein a subset of boxes underwent drought conditions, while the remaining boxes were regularly watered as a control group. The sequence transitions to (c), (d), and (e), acquired via a user camera, depicting broader perspectives of the entire box. (c) illustrates the growth stage of the root system, (d) portrays the plant during drought conditions, and (e) presents the rewatered plant. This collection of images offers a timeline of the experimental design and highlights the different stages of the study, enabling a better understanding of the temporal progression and experimental manipulations performed during the research. However, it is not representative of our imaging setup to acquire RGB and HSI data.

## 4. Applications

Our dataset has multiple applications. One of the first would be to utilize HS signatures to supplement existing root phenotypic traits with more in-depth physiological evaluation ([46], Root Phenotyping section). A couple studies have shown improved phenotyping prediction through added HS information [33, 34]. Researchers may also use the data to analyze root growth, architecture, and turnover of dense root systems [35, 36]. Some images contain other potential objects of interest such as fungus, mold, and algae which may be studied at their various timesteps to determine possible interactive dynamics between root and rhizosphere. By example, previous work has studied root-fungal relationships in peatland [37].

The additional HSI data can provide researchers with a more informative look at a plant’s health and physiology and may be applied to drought resiliency and nutrient concentration studies. By taking advantage of the dehydration and rehydration process in our dataset, researchers could predict plant water status in response to drought for two annual crop species [5, 6]. Creating additional links between HSI data and a plant’s health could enhance studies addressing micronutrient deficiencies in populations worldwide [47].

HyperPRI also presents ML experts with multiple challenges in root segmentation. Thin root features, with widths as narrow as 1 – 3 pixels, require robust algorithms for accurate identification and segmentation. Our data has a highly textured soil background that encourages exploration of texture analysis techniques. Finally, due to the high-resolution spectral data, reflectance differences between channels are reduced, meaning that chosen ML algorithms must be able to handle high correlation to be effective. While addressing the challenge of thin object features, ML experts can also contribute to solving segmentation problems with similar characteristics in other domains such as medical imaging [48].

## 5. Segmentation Methodology

We perform root segmentation on the subset of peanut roots using three approaches: spatial, spectral, and spatial-spectral. The first focuses on using only the RGB data and a UNET architecture [49] to segment the roots. The second utilizes a UNET-like autoencoder that has no knowledge of spatial neighborhoods within HSI data. The final method combines UNET segmentation and spectral signatures using a UNET with a 3D convolution.

### 5.1. Data Splits

Due to the low amount of data, we chose to evaluate models using a 5-split cross-validation. Within each split, data is split across boxes (ie. Different days for each box are kept as a group). In other words, if box 33 is in the training split, this is true for all images from Jun 24 to Aug 21. When deciding the splits for the peanut roots, it was thought that phenotype 1 (genotype TUF Runner 511) and phenotype 2 (genotype 10×34-4-4-1-2) would need approximately equivalent training-validation representation to prevent bias in learned features. To understand the peanut subset, we trained a UNET on a portion of both phenotypes (20 and 19 images, respectively) such that they both have approximately equivalent timestep representation. When training UNET from scratch for five random initializations on phenotype 1 (phenotype 2) and evaluating on phenotype 2 (phenotype 1), the resulting DICE, root intersection-over-union (+IOU), and average precision (AP) values do not demonstrate statistically significant differences between the two (Table 2 and Fig. 3a). The p-values for all three were computed with a 2-sided unpaired, unequal variance Student t-test using Excel.

**Figure 3:**
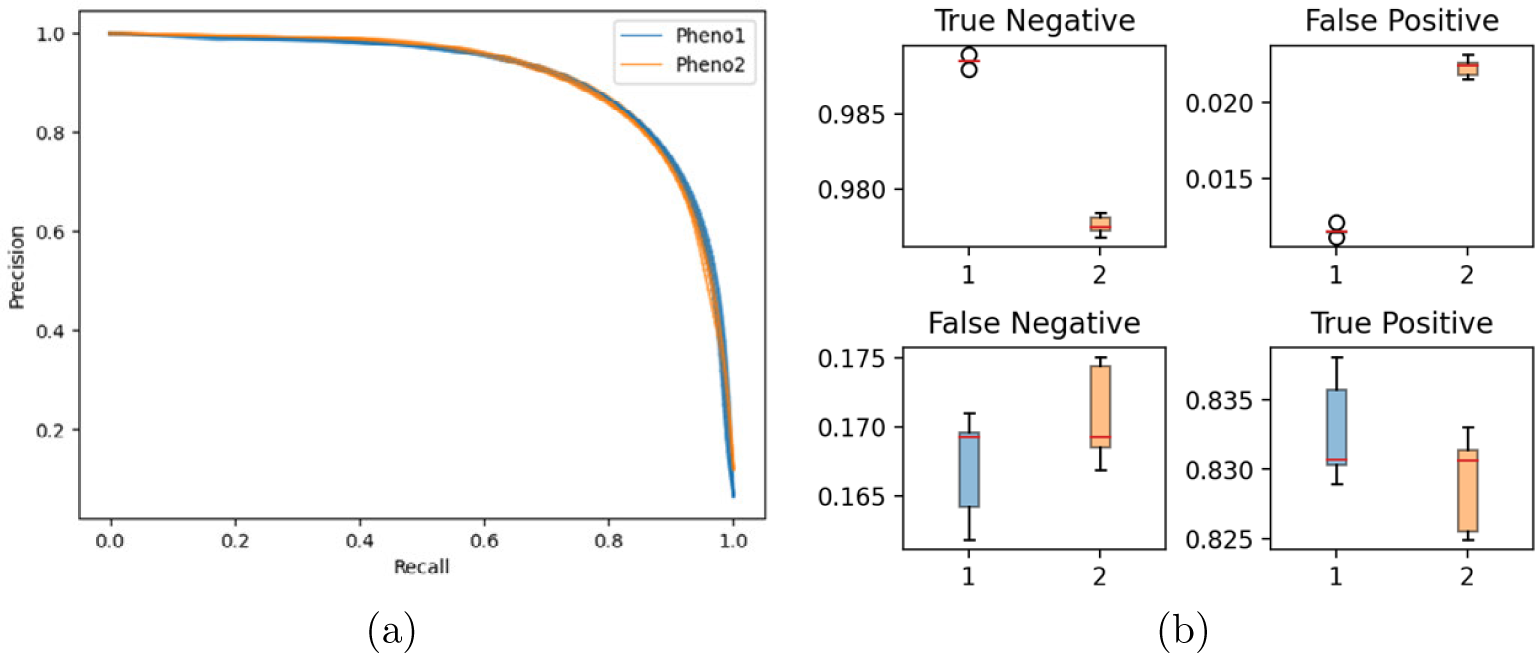
Precision-Recall curve (a) and confusion matrix statistics (b) for UNET trained on phenotype 1 and phenotype 2. In (a), Pheno1 and Pheno2 correspond to training on phenotype 1 and 2, resp. For the confusion matrix statistics, X-axis labels correspond to which phenotype was used for training data, and red lines show median results across all splits.

**Table 2:** Mean, deviation, and max (in parentheses) performance for training a UNET on Phenotype 1 and Phenotype 2. Results are for validating on Phenotype 2 and Phenotype 1, respectively.

|  | Tr on Phenotype 1 | Tr on Phenotype 2 | FDS | p-value |
| --- | --- | --- | --- | --- |
| **DICE** | $0.835 \pm 0.001$ (0.837) | $0.833 \pm 0.003$ (0.837) | 0.0010 | 0.167 |
| **+IOU** | $0.717 \pm 0.002$ (0.720) | $0.713 \pm 0.004$ (0.720) | 0.0027 | 0.163 |
| **AP** | $0.909 \pm 0.001$ (0.913) | $0.910 \pm 0.002$ (0.913) | 0.0002 | 0.366 |

Fig. 3b shows normalized confusion matrix statistics when using the best threshold per random seed. The normalizing term per row is each image’s number of true root or soil pixels (ie. normalize TN and FP with TN+FP).

The boxplots show much less variance across TN and FP samples because the root-soil pixel ratio for this subset of phenotype 1 and 2 images were 0.139 and 0.073, respectively. Fig. 3b also shows the median TP results are the same, adding further evidence that the UNET tends to learn similar features between the two phenotypes.

In addition to the above phenotype comparisons, the Fisher Linear Discriminant Score (FDS) [50]

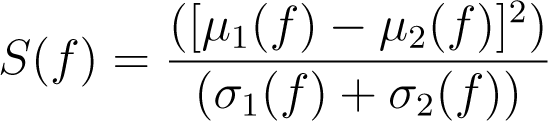

is used as a scoring function to quantify the distance between means and deviations of two distributions. *µ* and *σ* are the mean and standard deviation of DICE, +IOU, and AP results per phenotype (represented by *f*). Table 2 shows that – compared to the difference between root and soil pixels in Fig. 7 – the difference between training results for the two phenotypes is low. A couple other ways to compare the two phenotypes are by counting the proportion of certain edge types with edge histogram descriptors (EHDs) [51] in Fig. 4 and comparing either the root or soil pixel spectra (Appendix A, Fig. A.15). In both cases, the computed FDS between phenotype 1 and 2 characteristics remain on the same order as scores in Table 2. Thus, moving forward, we do not maintain equivalent phenotypic representation in a 5-split organization of the dataset but do maintain an approximately equivalent training-validation ratio. In total, there were 59 images used, and training had between 43 and 45 images for each split (Table A.6). Lastly, for our test set, we use two images from rhizobox 40 at days 2022-08-15 and 2022-08-24 that are drought-level (‘dry’) and rewatered (‘wet’), respectively.

**Figure 4:**
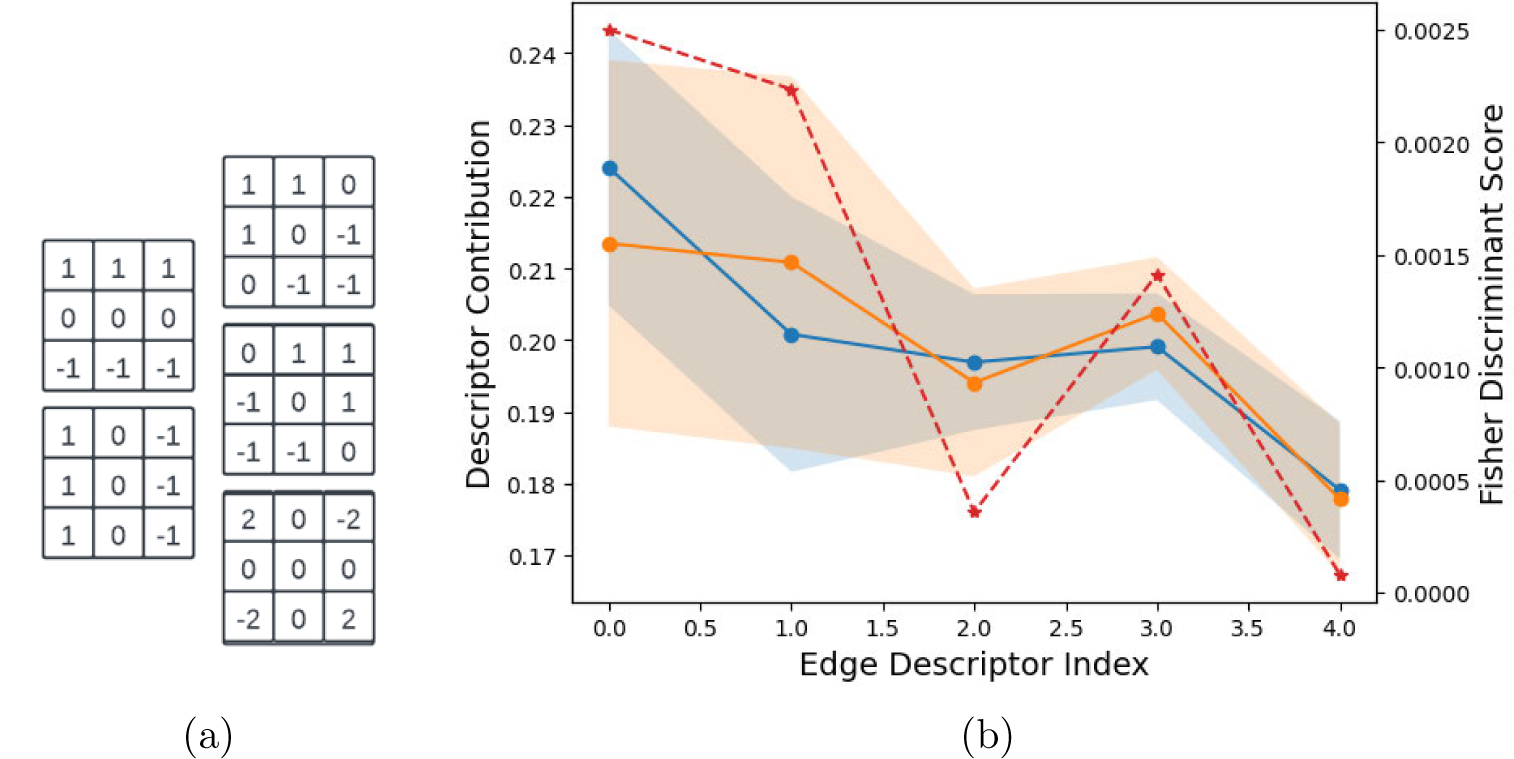
Edge Histogram Descriptor comparison between phenotype 1 (blue) and 2 (orange). The edge filters are shown in (a), with index increasing from top-to-bottom, left-to-right. In (b), the measured difference between each EHD bin (red, dashed) demonstrates that the difference between proportions of different edge types is greatest for the horizontal and vertical edges (index 0 and 1).

### 5.2. Spectral Characteristics of Viewing Pane

At each stage of data acquisition, the Lexan™ viewing pane is not removed, so another factor to investigate when considering hyperspectral features is the plastic sheet’s effect on the spectral signatures. Using a rhizobox of the same make as above and only filled with Turface soil, we imaged the box in dry and rewatered scenarios with the clear plastic sheet in place and removed. Summary statistics across the entire image are shown in Fig. 5. We see that the clear plastic sheet absorbs or prevents a proportional amount of reflectance for both scenarios of dry and wet soil. While the signatures’ magnitudes change, the shape of the signatures hardly change when the sheet is removed, and this change becomes almost imperceptible when the soil is wet. To further quantify this statement, we use the spectral angle difference to compare mean spectral signatures of the images with and without Lexan. They are respectively represented by **x** and **y**. For dry and wet boxes, the spectral angles were found to be 0.0497 rad (2.8477°) and 0.0220 rad (1.2628°), resp. The result shows that both mean spectral signatures are similar vectors and separated in value only by scaling differences. The sheet is therefore left in place when acquiring data at each stage of the aforementioned timeline (Section 3).

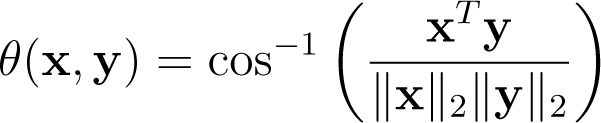

**Figure 5:**
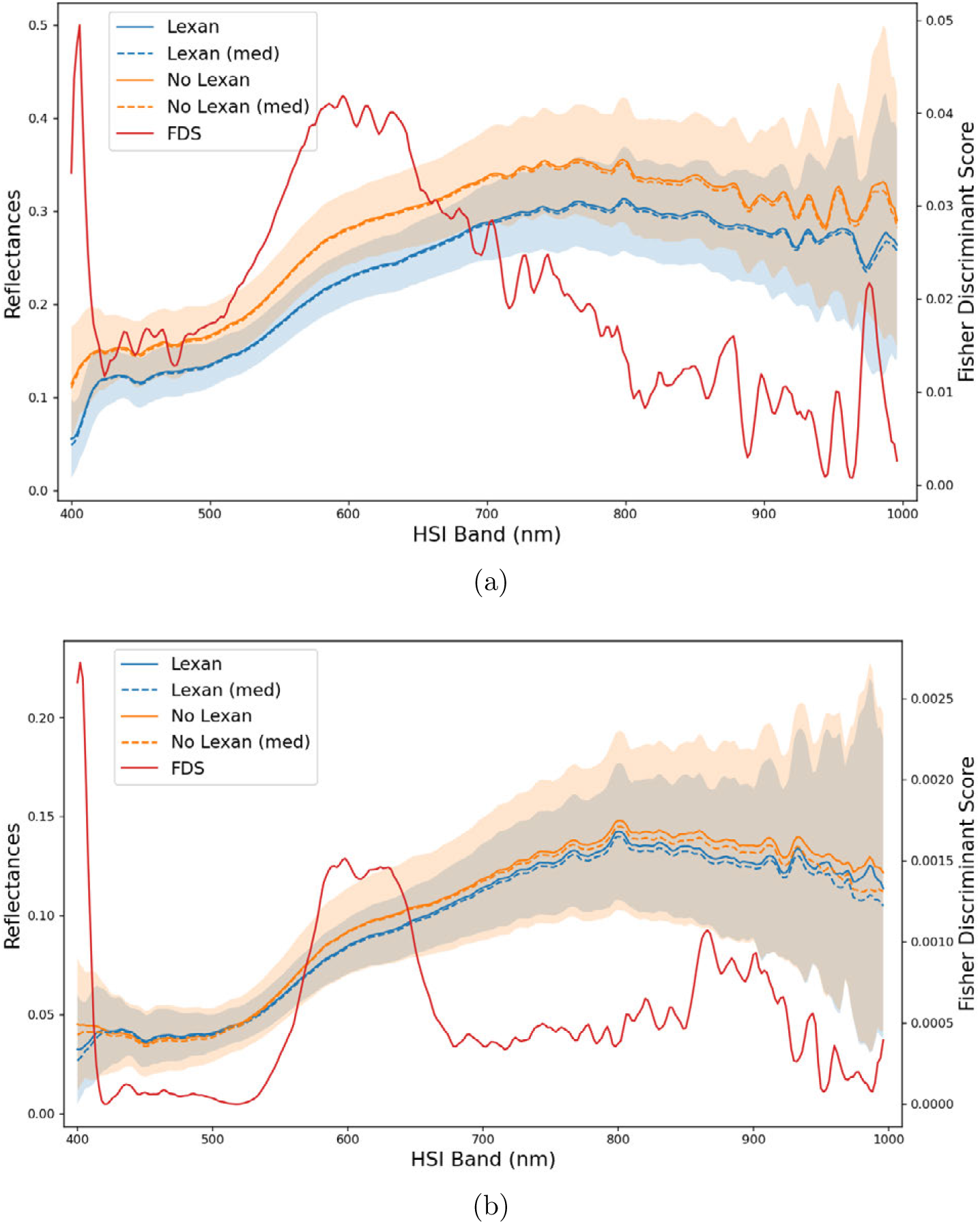
Comparison of summarized spectral reflectances for a rhizobox filled with (a) dry and (b) wet Turface soil when the Lexan™ is in place or removed. Regarding summary reflectance statistics, the solid lines, dashed lines, and shading between the lines represent means, medians, and standard deviations, resp. (a) For the dry soil, the plastic sheet absorbs or prevents a proportional amount of reflectance. (b) For the wet soil, the difference in reflectance drops by a factor of approximately 20. Both scenarios demonstrate a spectral signature that remains the same regardless of whether the plastic sheet is attached.

### 5.3. Model Architectures

Once the data is divided into five splits (Table A.6), each split is input to three different UNET models. The models use either conventional RGB image analyses, only HS signatures, or conventional image analyses coupled with HS signatures.

First, the “spatial” method does conventional image analysis using RGB images. A UNET model of depth 5 is used with BatchNorm [52] in the double convolution feedforward stages. The filter sizes and number of feature maps per each encoding depth is the same as the original work [49] but with paddings of 1 on either side. To train from randomly initialized weights, we use the Adam optimizer [53] with default parameters (0.99, 0.999, 0.001) and no weight decay. Models are trained for a maximum of 1000 epochs with early stopping after 250 epochs of no binary cross-entropy loss improvement. Only weights demonstrating better validation DICE (threshold 0.5) are saved for evaluation.

Second, the “spectral” method uses only HS signatures. A UNET-like architecture (named SpectralUNET) is trained on pixels’ spectral signatures that have D bands. To achieve a comparably-sized network, SpectralUNET’s encoder has layers of size 1650, and its decoder has layers of size 1650 that use concatenated information using skip connections from the encoder layers as inputs. Its final output layer is a single neuron that outputs the model’s confidence that a pixel is root. Each layer input and hidden layer is a sequence of a linear layer, ReLU nonlinearity, and BatchNorm layer. A figure of the architecture is provided in Fig. 6. The same optimizer and number of training epochs remain the same as in the spatial method.

**Figure 6:**
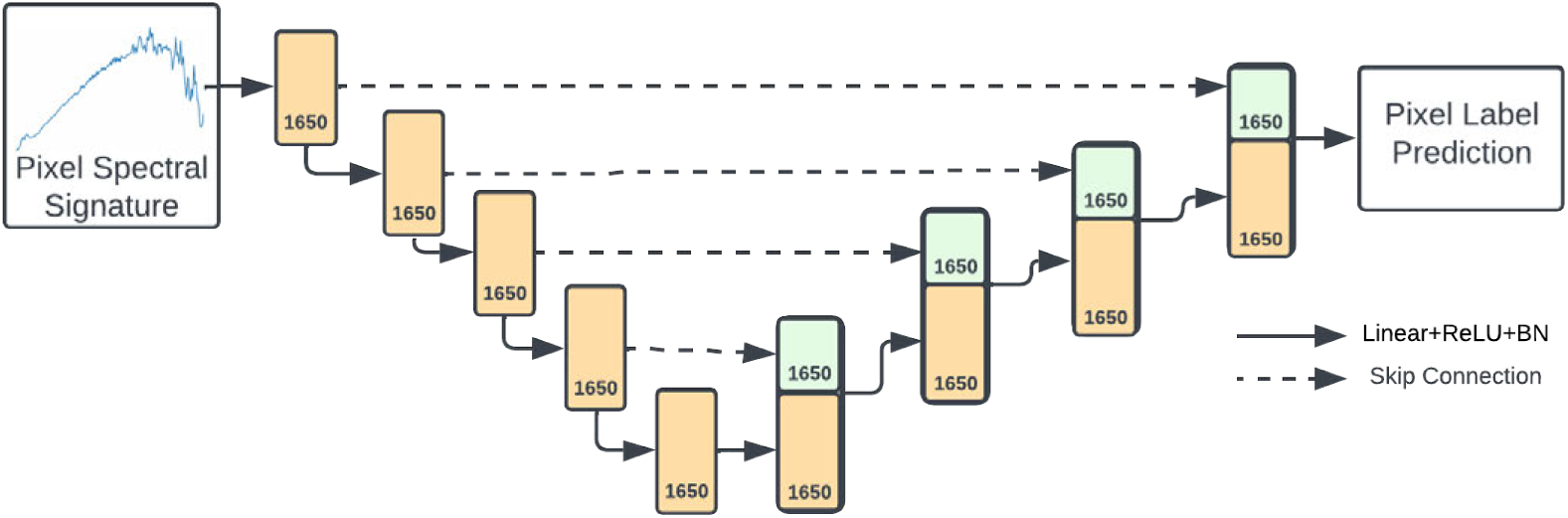
SpectralUNET architecture. The numbers represent the dimensionality of a layer’s output. Blocks in green represent concatenated information from previous feedforward layers.

**Figure 7:**
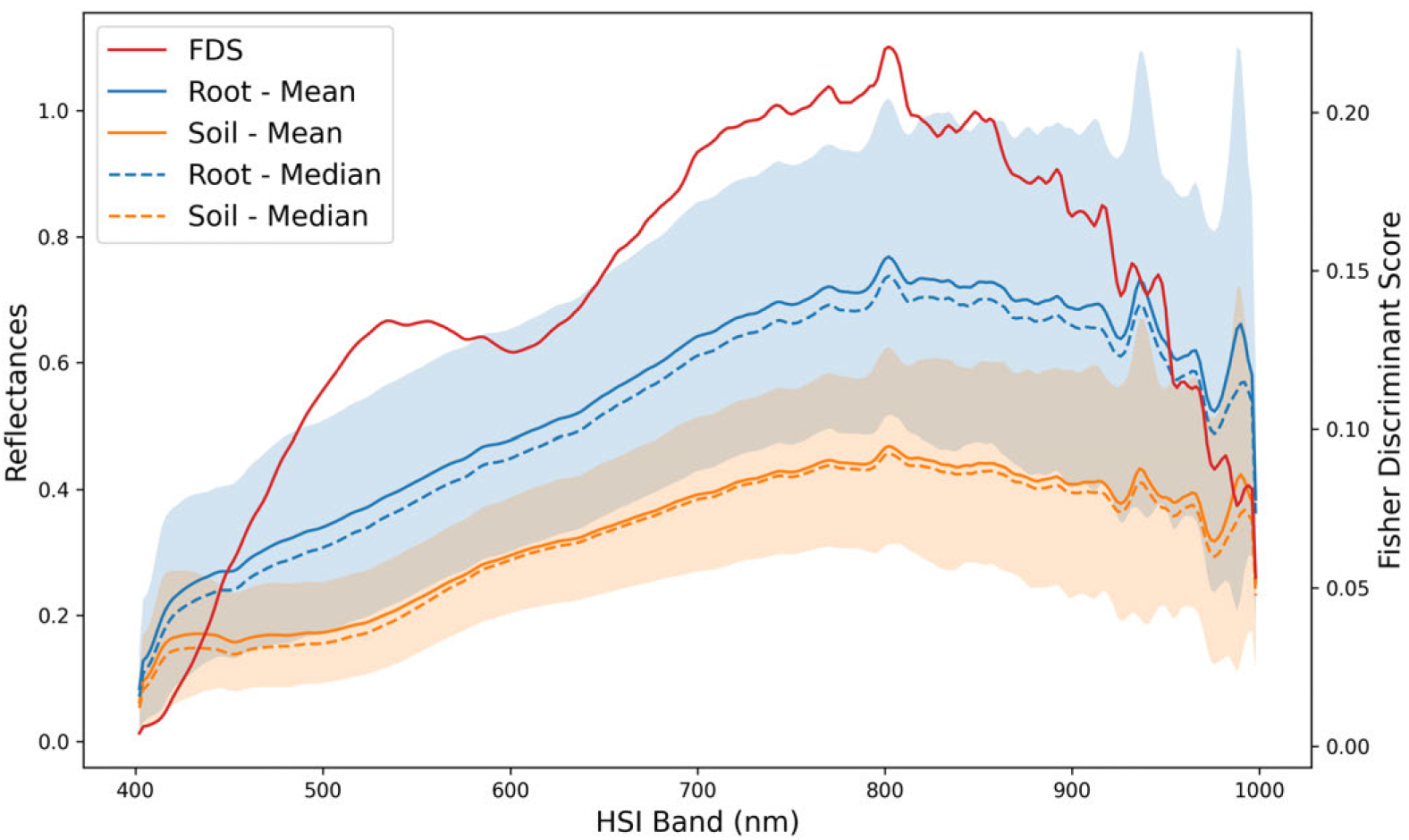
Reflectance comparison between root and soil spectral signatures across 15 peanut rhizoboxes. Shading in the respective colors represents a single standard deviation from the solid lines. Fisher’s linear discriminant score is computed at each band. The means and deviations were computed after smoothing each signature with a moving average of size 3.

Third, combining conventional image analysis with HS signatures to create a “spatial-spectral” method, we use a UNET whose first set of layers is instead a 3D convolutional layer that uses a filter of size D*×*3*×*3 (CubeNET) and goes to 64 channels as in the spatial method. Following this first convolution, the order and size for the rest of the convolutional layers is the same as the spatial method. We use the same Adam optimizer configuration as the spatial method with default parameters and 1000 epochs of training with early stopping of 250 based on the validation BCE loss. Compared to keeping all *D* channels, we chose to learn weights to 64 channels to keep the model size (ie. degrees of freedom) comparable between CubeNET, UNET, and SpectralUNET. Thus, any performance increases are less (if at all) attributed to an increase in the number of trainable model parameters.

To minimize model overfitting due to the small dataset, we saved model parameter checkpoints at each epoch when a model improved upon its validation BCE loss. These parameter checkpoints are reloaded to get results and segmentation predictions in subsequent sections.

## 6. Segmentation Results

Here, we discuss low and high HS band choices, segmentation performance, limitations for the data and learning models, and future work.

### 6.1. HSI Data Preprocessing

Prior to using the spectral or spatial-spectral methods, we must consider what region of the spectral signature to use as input. Therefore, we consider the mean and standard deviation for 33% of the root and soil pixels from 15 peanut rhizoboxes. From Jun 24th to Aug 8th, the spectral signatures are compared as in Fig. 7 and 8. Fig. 7 shows that root pixels have signatures with higher average reflectance but also greater variance than soil. Also, Fig. 8 shows that the distribution of both signatures have skewed statistics with longer tails toward higher reflectance. We use the FDS to measure the separability of the root and soil distributions. Scores are computed for the reflectance means and standard deviations for root and soil pixels at the each given wavelength across all 15 boxes. As shown in Fig. 7, the discriminant score for the two classes increases significantly by 450 nm and drops sharply after 925 nm. With this in mind, we choose to train and evaluate the spectral/spatial-spectral methods with spectral information from 450 – 926 nm (approximately cube index 25 to 263, exclusive) of each HSI cube. Therefore, the input for the SpectralUNET and CubeNET are 588, 544 *×* 238 and 238 *×* 608 *×* 968, respectively.

**Figure 8:**
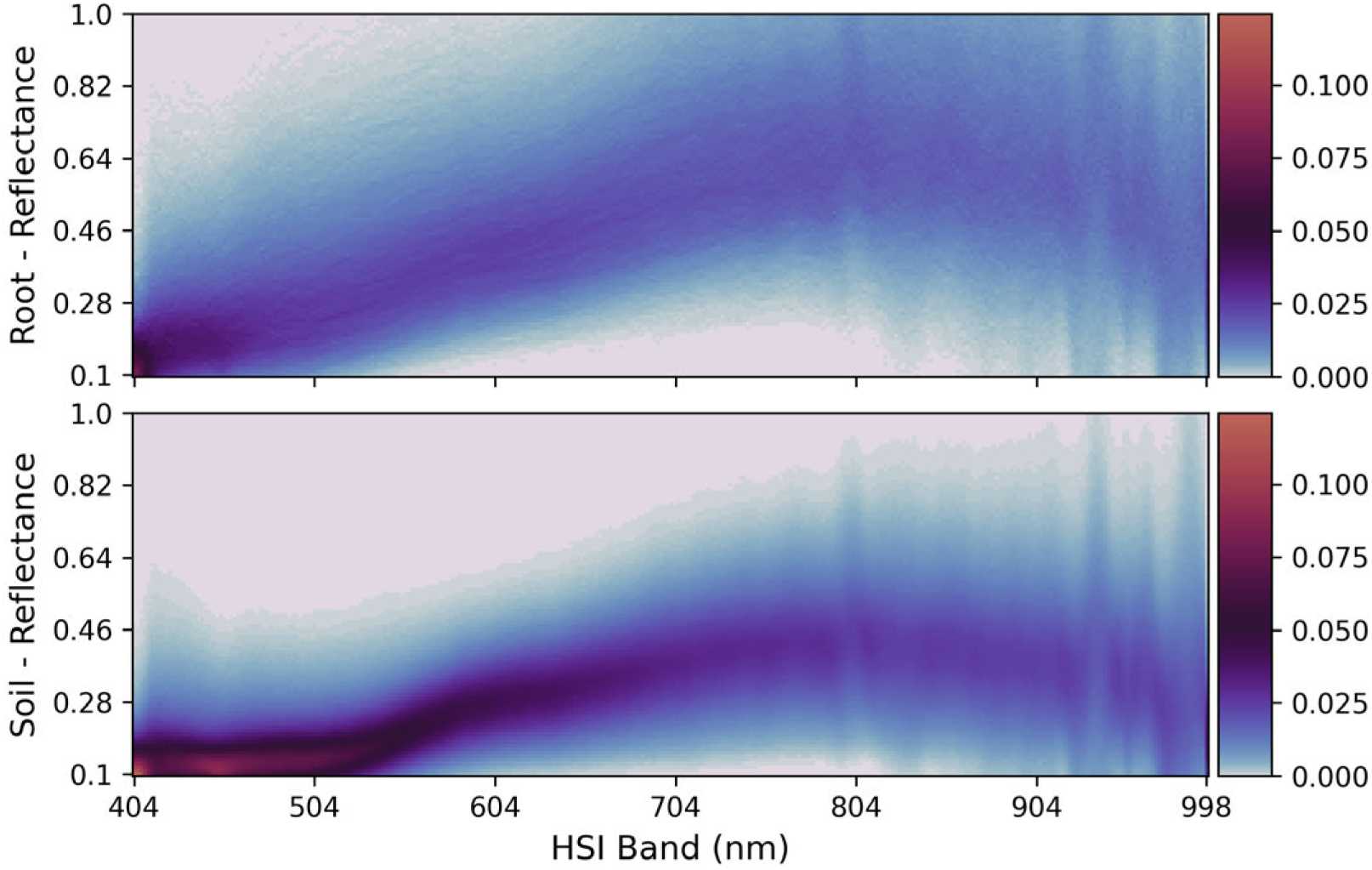
Raw reflectance comparisons between root and soil spectral signatures across 15 peanut rhizoboxes. Both root and soil plots show their distributions are skewed toward a higher reflectance compared to their means.

### 6.2. Model Comparisons – Validation Data

Final training loss, number of trainable weights, and quantitative validation results for UNET, SpectralUNET, and CubeNET are shown in Tables 3 and 4, respectively. To produce Table 3, the thresholds per each model are chosen according to their best validation DICE score on each split. In this way, we are comparing the models’ “best” expected performance when used for annotating unseen data (ie. test data). UNET continues to perform well when focusing on spatial information in the visible RGB channels. Due to inherent noise in the spectral reflectance (Fig. 7 and 8), SpectralUNET performs the worst. Across all five splits, the CubeNET has better performance and adds a little under one second of per-image CPU inference time (Table 3). We also note that including spectral information with spatial methods decreases the variation in DICE results for CubeNET compared to the other two.

**Table 3:** Number of trainable weights, CPU-based inference time, and 5-split validation Binary Cross-Entropy loss for UNET, SpectralUNET, and CubeNET. Lower BCE Loss is better.

|  | Val BCE Loss | # Weights | Per-Image Time (s) |
| --- | --- | --- | --- |
| **UNET** | $0.080 \pm 0.015$ | 31,043,521 | $4.633 \pm 0.065$ |
| **SpectralUNET** | $0.146 \pm 0.022$ | 30,388,051 | $292.815 \pm 0.291$ |
| **CubeNET** | $0.077 \pm 0.014$ | 31,178,881 | $5.425 \pm 0.224$ |

**Table 4:** Quantitative validation results for UNET, SpectralUNET, and CubeNET. Best results across all models are in bold. Higher DICE, +IOU, and AP scores are better.

| Split | UNET |  |  | SpectralUNET |  |  | CubeNET |  |  |
| --- | --- | --- | --- | --- | --- | --- | --- | --- | --- |
|  | DICE | +IOU | AP | DICE | +IOU | AP | DICE | +IOU | AP |
| 1 | 0.830 | 0.709 | 0.915 | 0.663 | 0.496 | 0.719 | 0.841 | 0.725 | 0.923 |
| 2 | 0.825 | 0.702 | 0.909 | 0.720 | 0.563 | 0.793 | 0.831 | 0.711 | 0.915 |
| 3 | 0.864 | 0.761 | 0.942 | 0.767 | 0.622 | 0.829 | 0.868 | 0.766 | 0.946 |
| 4 | 0.843 | 0.729 | 0.922 | 0.764 | 0.618 | 0.832 | 0.843 | 0.729 | 0.918 |
| 5 | 0.827 | 0.705 | 0.906 | 0.673 | 0.507 | 0.731 | 0.837 | 0.720 | 0.914 |
| **Mean** | 0.838 | 0.721 | 0.919 | 0.717 | 0.561 | 0.781 | **0.844** | **0.730** | **0.923** |
| **SD** | 0.015 | 0.022 | 0.013 | 0.044 | 0.053 | 0.048 | **0.013** | **0.019** | **0.012** |

Fig. 9 compares the precision-recall curves between methods and shows that CubeNET handles the precision-recall tradeoff better than the other methods and may perform the best on unseen examples. UNET has slightly more variance in its performance but performs well, and SpectralUNET shows the most variance and the worst performance. Combining UNET’s spatial filters with the high spectral variance seen in Fig. 8 led to more consistent and generalizable results. Similar conclusions appear to be found in Fig. 10; the main difference between UNET and CubeNET is in the improvement of TP’s mean and consistency (distribution tightness).

**Figure 9:**
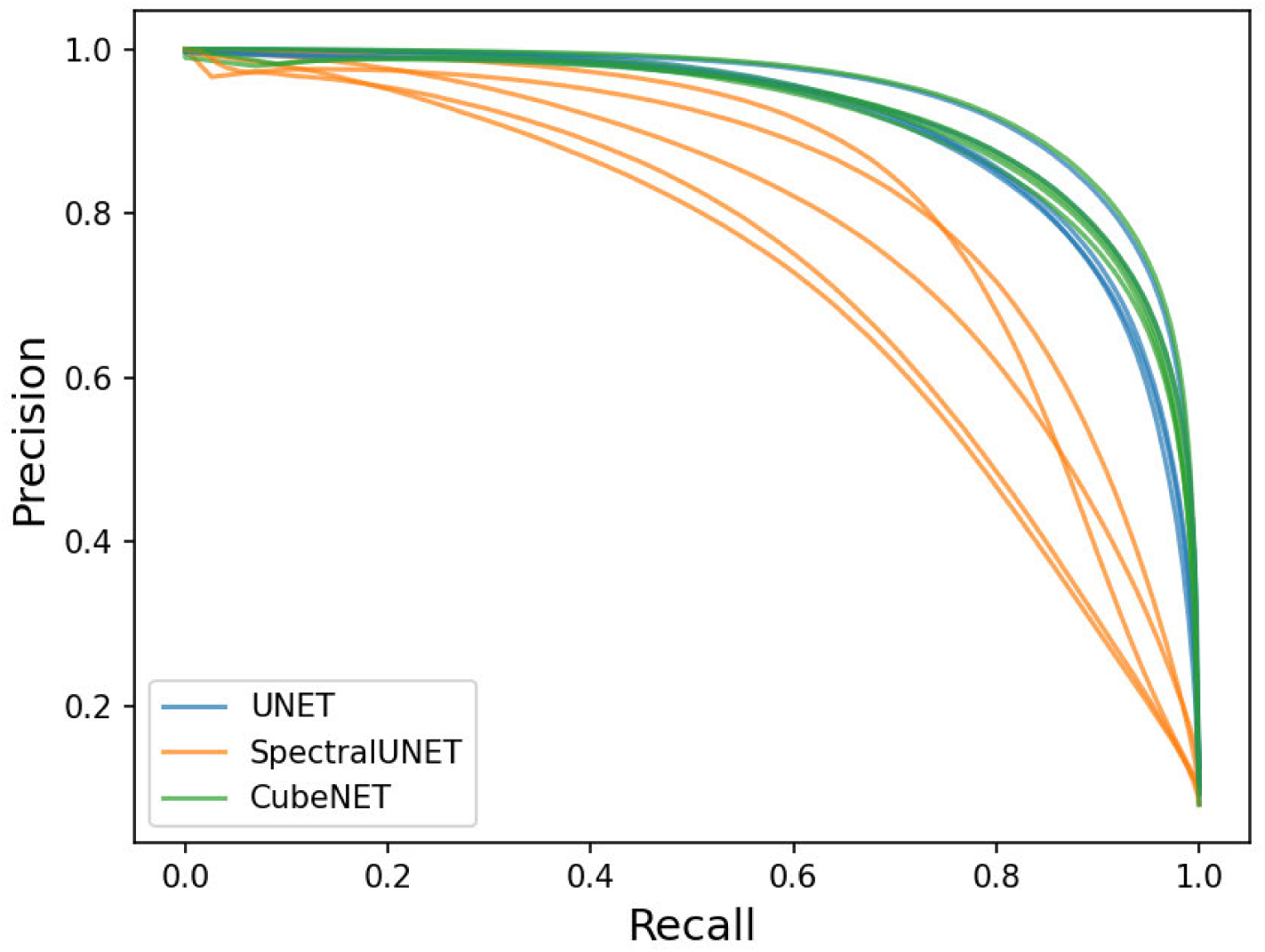
Precision-Recall curves for UNET, SpectralUNET, and CubeNET on the five splits of data. Across multiple confidence thresholds, CubeNET demonstrates slightly more robustness.

**Figure 10:**
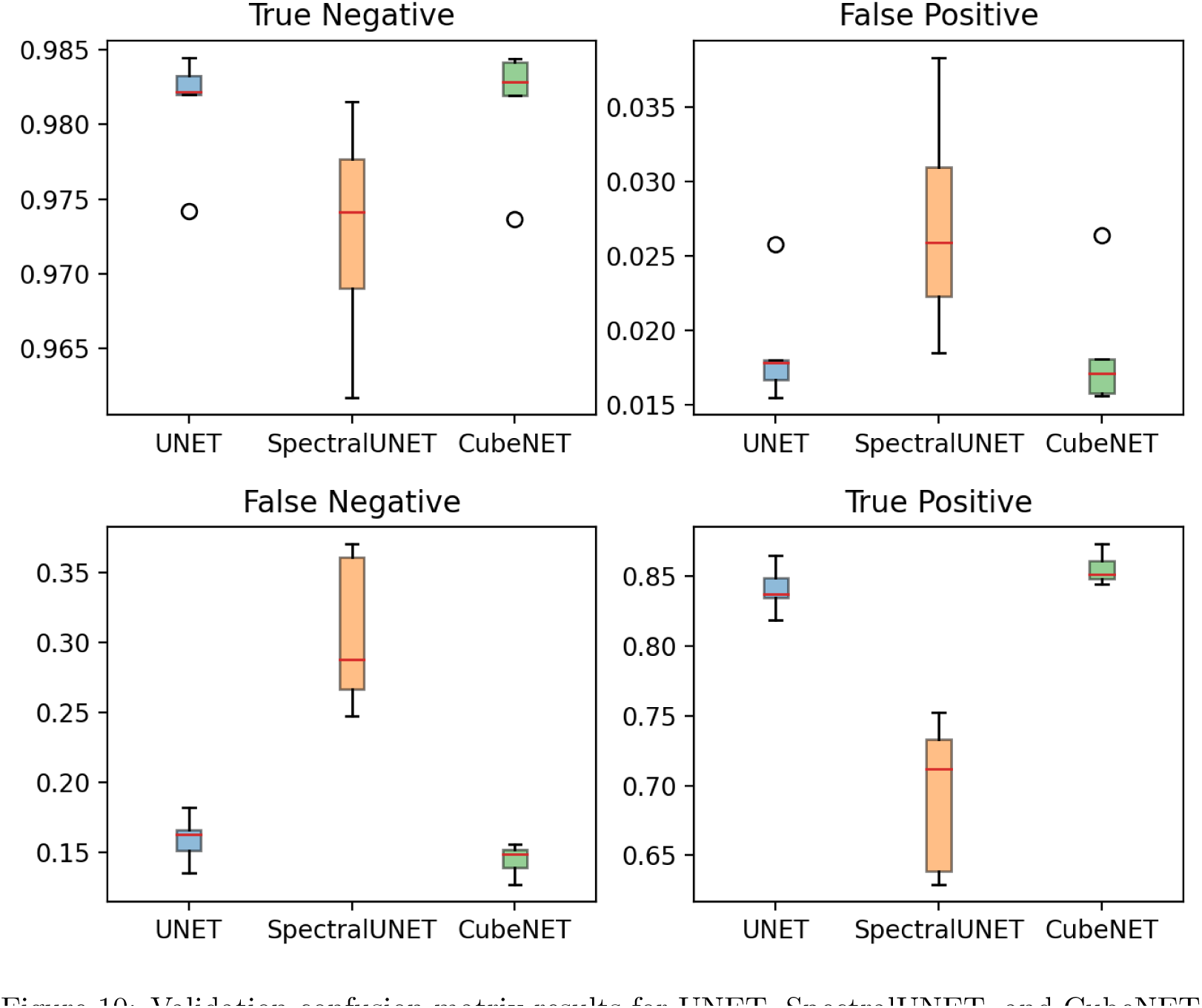
Validation confusion matrix results for UNET, SpectralUNET, and CubeNET across the five splits of data. Utilizing both hyperspectral and spatial features, CubeNET achieves improved precision and greater consistency in predicting roots.

When comparing segmentation masks as seen in Fig. 11, CubeNET segments roots in dry images and thin roots more consistently than UNET, but sometimes the peanut pegs are not fully identified along with the root class (see second row of Fig. 11). Overall, SpectralUNET typically has more FP noise than the other two, especially when the soil’s visible spectrum is like roots. However, all three tend to show some overstepping of root boundaries and could benefit from post-processing to clean up results. Additional selected segmentation results for all three models may be seen in Appendix B (Fig. B.16, B.17, B.18).

**Figure 11:**
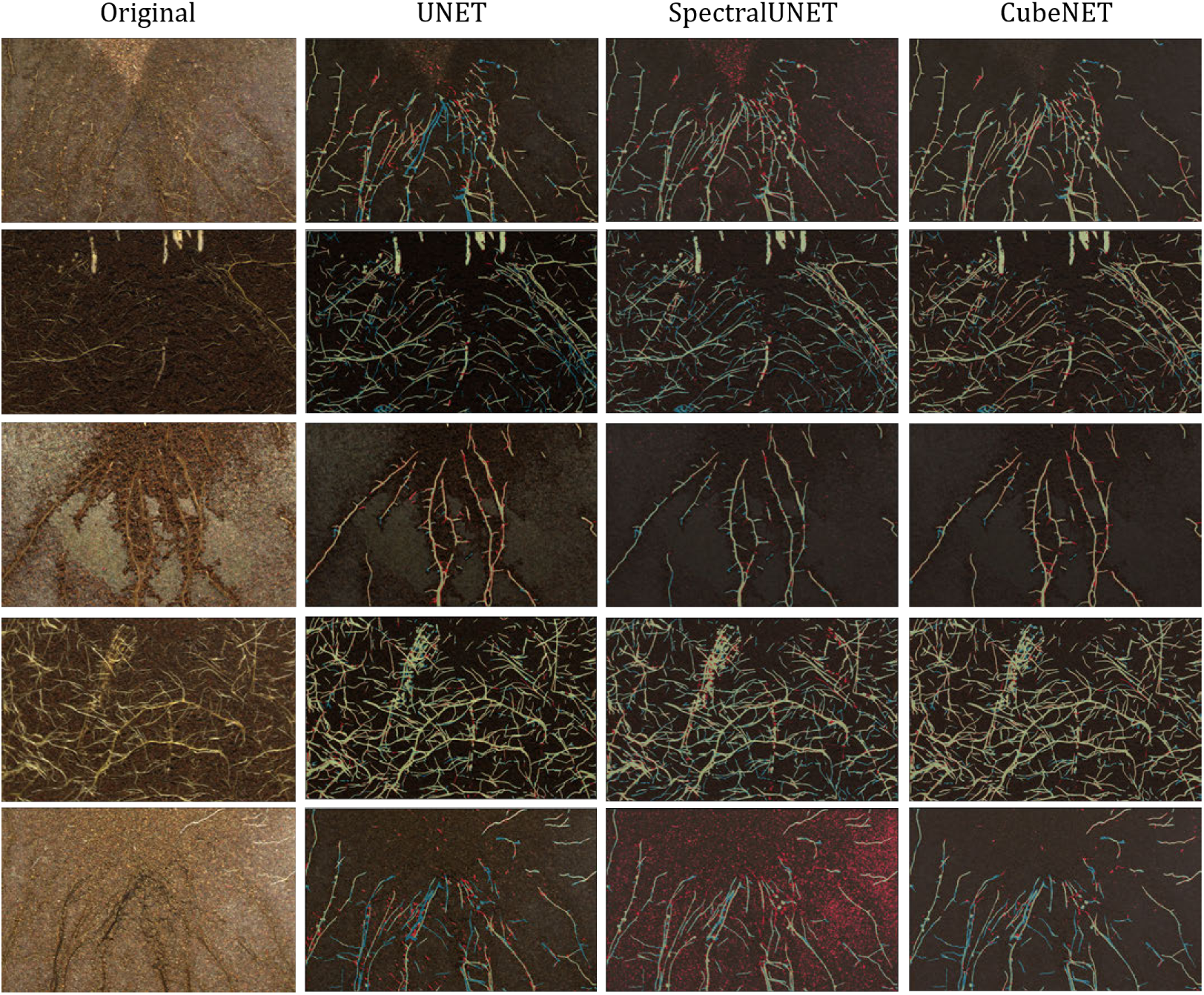
Selected segmentation results for UNET, SpectralUNET, and CubeNET. Each row corresponds to one split of data (ie. the second row takes segmentations from split 2). Darkened image pixels are true negatives. Red pixels are false positives. Blue pixels are false negatives. Yellow-green pixels are true positives.

From the results, we may conclude that deep learning models attempting to segment roots from soil must utilize spatial information. We may also conclude that - although subpar alone - spectral signatures combined with spatial information can cause models to learn a better, more generalized understanding of target objects when solving the problem of semantic segmentation.

### 6.3. Model Comparisons – Test Data

Using the unseen dry and wet box data for rhizobox 40, we do a final comparison of the three models. To construct the results seen in Table 5 and Figures 12 and 13, we used the saved models and cross-validated thresholds from training on all five splits.

**Figure 12:**
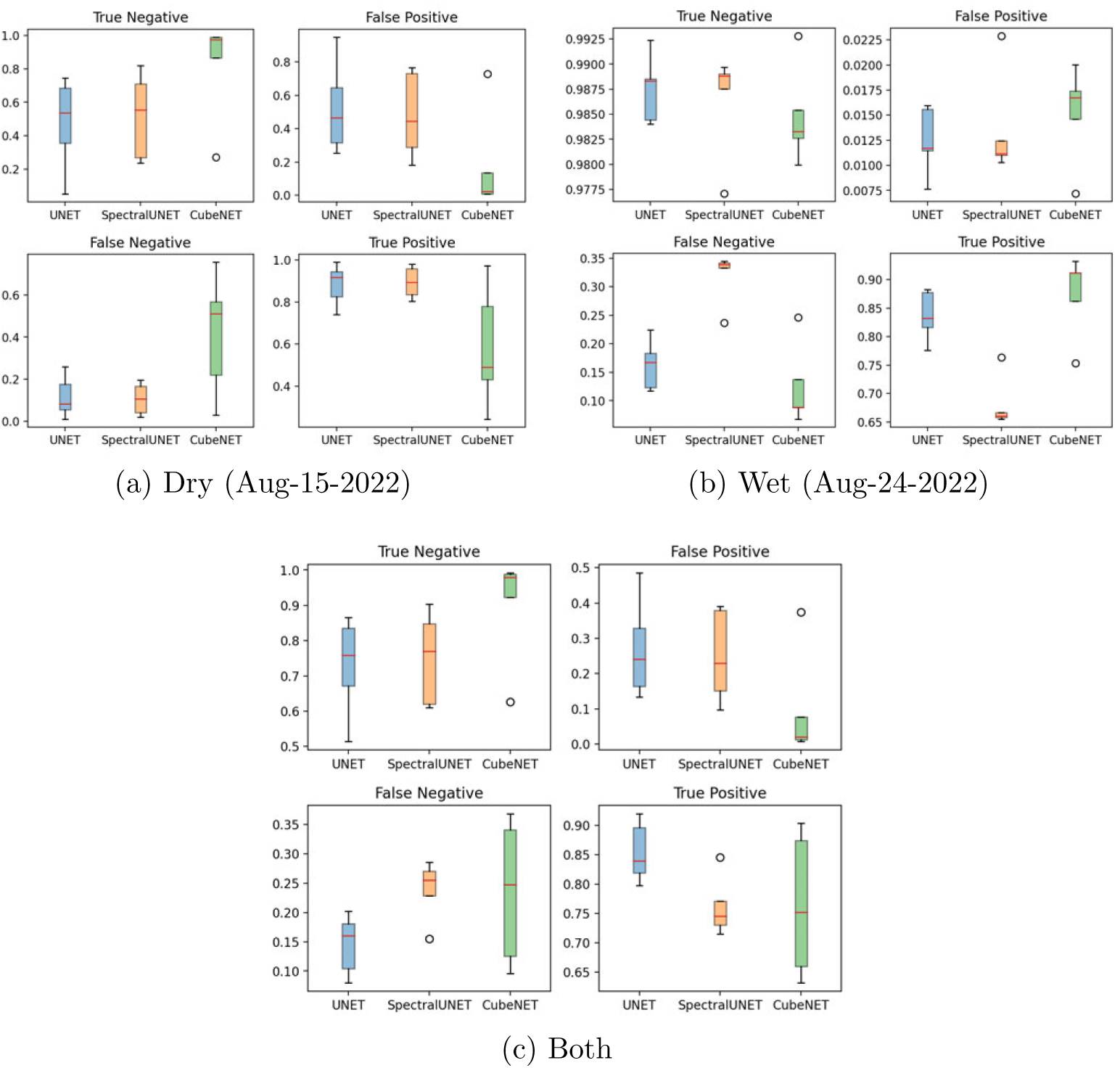
Test confusion matrix results for UNET, SpectralUNET, and CubeNET. Using both hyperspectral and spatial features, CubeNET undersegments while both UNET and SpectralUNET tend to oversegment.

**Figure 13:**
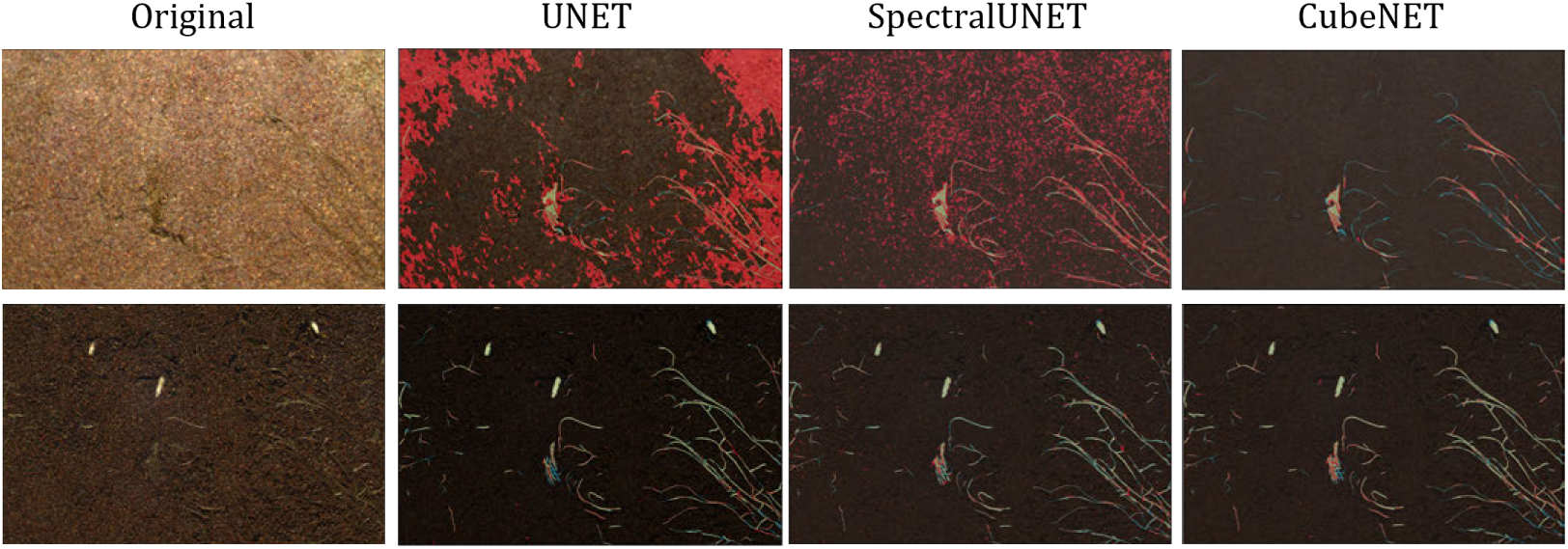
Test segmentation results for UNET, SpectralUNET, and CubeNET. The rows correspond to dry and wet box images, resp. Darkened image pixels are true negatives. Red pixels are false positives. Blue pixels are false negatives. Yellow-green pixels are true positives.

**Table 5:** Quantitative test results for UNET, SpectralUNET, and CubeNET. Best results across all models are in bold. Higher is better for all metrics.

| Model |  | Dry | Wet | Both |
| --- | --- | --- | --- | --- |
| UNET | Acc | $0.482 \pm 0.245$ | **$0.983 \pm 0.001$** | $0.733 \pm 0.123$ |
| | DICE | $0.073 \pm 0.025$ | **$0.757 \pm 0.010$** | $0.162 \pm 0.053$ |
| | +IOU | $0.038 \pm 0.014$ | **$0.609 \pm 0.013$** | $0.089 \pm 0.031$ |
| | AP | $0.138 \pm 0.049$ | $0.836 \pm 0.007$ | $0.226 \pm 0.079$ |
| SpectralUNET | Acc | $0.525 \pm 0.227$ | $0.976 \pm 0.003$ | $0.751 \pm 0.114$ |
| | DICE | $0.084 \pm 0.037$ | $0.653 \pm 0.016$ | $0.161 \pm 0.064$ |
| | +IOU | $0.044 \pm 0.020$ | $0.485 \pm 0.017$ | $0.089 \pm 0.039$ |
| | AP | $0.201 \pm 0.082$ | $0.697 \pm 0.011$ | $0.220 \pm 0.083$ |
| CubeNET | Acc | **$0.815 \pm 0.268$** | $0.981 \pm 0.002$ | **$0.898 \pm 0.134$** |
| | DICE | **$0.273 \pm 0.145$** | $0.751 \pm 0.010$ | **$0.471 \pm 0.206$** |
| | +IOU | **$0.166 \pm 0.097$** | $0.601 \pm 0.013$ | **$0.329 \pm 0.163$** |
|  | AP | **$0.345 \pm 0.079$** | **$0.843 \pm 0.009$** | **$0.610 \pm 0.109$** |

Table 5 shows the mean and standard deviation of metrics for all three models, and Figure 12 shows the spread of the confusion matrix results for the dry and wet images. Overall, the summarized results show that CubeNET outperforms UNET and SpectralUNET on test data. The difference in performance is even greater when considering segmentation results on a dry image; CubeNET is shown to undersegment the roots while the other two oversegment and have two or three times more false positives (see also Fig. 13). We may find a suggested reason for this when comparing UNET and SpectralUNET. On the dry image, SpectralUNET outperforms UNET on average across all metrics. This suggests that CubeNET’s performance boost primarily comes from incorporating hyperspectral information. This further suggests that future studies which look at in situ roots under drought-level conditions should also use HSI data to differentiate roots and soil.

### 6.4. Limitations

Although sufficient for semantic segmentation purposes, the fully-annotated dataset has certain limitations to consider. First, the full set of annotations was completed by multiple co-authors. By way of demonstration, Fig. 14 shows examples of overlapped ground truth masks for four selected images from the two highest contributing annotators. Both annotators agree on most of the root pixels. However, both annotators add thin roots or root edges they believe are in the image that the other does not. Quantitatively, we may compare the discrepancy in annotation through the sum of a logical XOR image normalized by the sum of the logical OR image. When we consider the four images in Fig. 14, from left to right, top to bottom, the fraction of discrepancy is 0.344, 0.219, 0.270, and 0.397 for an average of 0.308. A couple of the images were annotated with the VIA tool, leading to thicker annotations. Both annotators later used Photoshop or GIMP and drew thinner roots in later masks.

**Figure 14:**
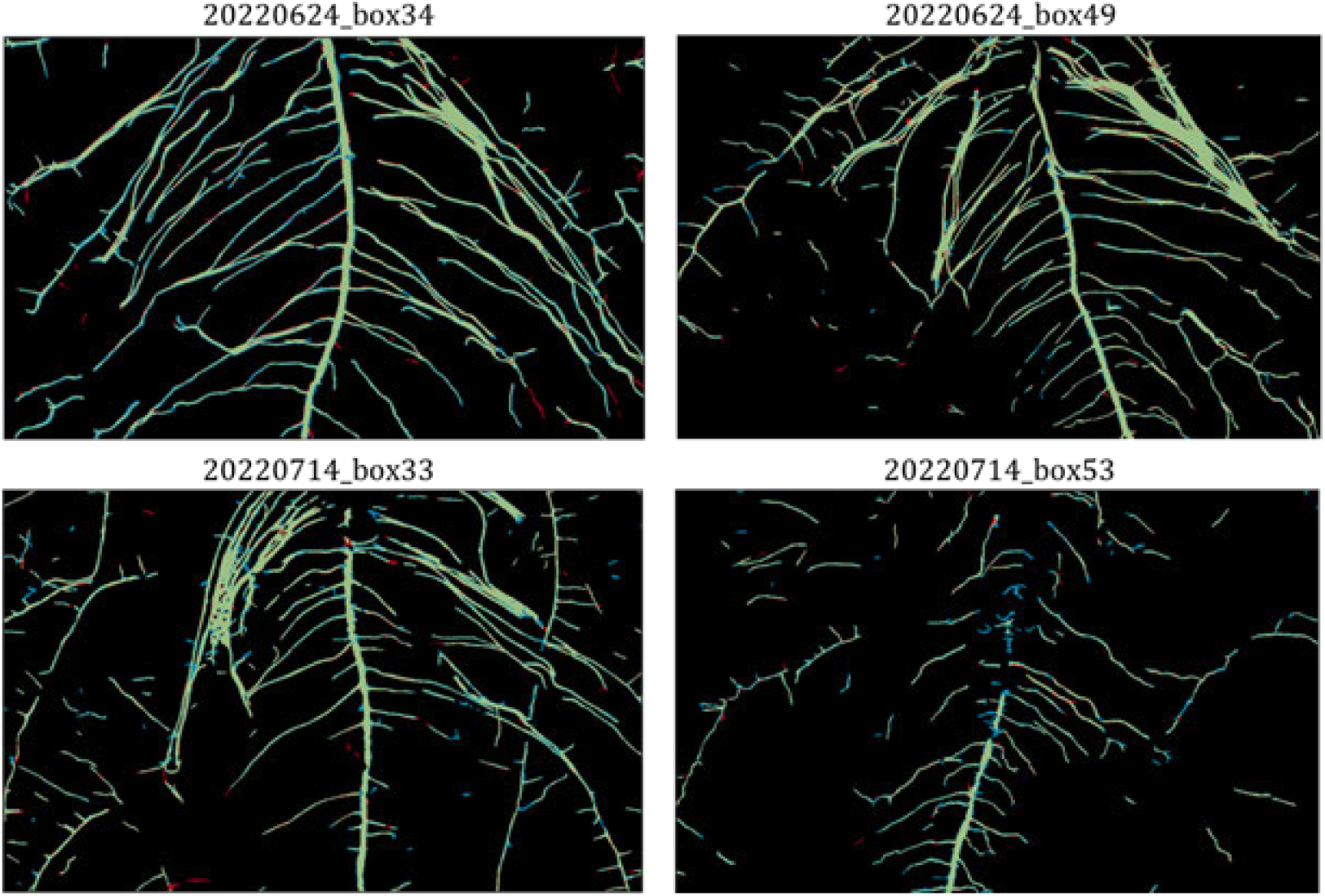
Selected dataset annotation comparisons for two annotators’ ground truth masks. Darkened and green pixels represent consistent annotations between Person 1 and 2, while red and blue show differing true positive annotations for Person 1 and 2, respectively.

Second, the root and soil objects often became indistinguishable depending on the box water conditions. Dry soil often had the same RGB colors as living peanut roots, and wet soil often had the same RGB colors as dead peanut roots. In certain instances, dry/wet soil may be differentiated from living/dead roots through certain HSI bands. Additionally, some root annotations may be missing because the annotators may believe a thin object to be a scratch on the box instead of a new, thin root growing in the soil.

Although we could use models trained on other datasets or previously annotated HyperPRI data, the pretrained model and methods like RootGraph [54] would require manual checks of the masks. The segmentation checks take as much time as manual annotations from a blank mask. Alternative tools, such as RootPainter [55], may help with annotations but would likely require the finetuning of multiple models due to the high variability we see in our dataset’s soil conditions. The third limitation of the dataset is that the camera hardware tended to have increased noise toward the longer wavelengths, and approximately the last 35 bands (70 nm) most likely could not be used due to the excessive noise. As previously illustrated, Fig. 7 shows that the discriminant score decreased with increases in the signature variances. Lastly, some unexpected data features such as mold and algae appear in later peanut images and were at times difficult to distinguish from roots. Regarding the segmentation methods, CubeNET demonstrates clear but minimal improvements compared to UNET. Compared to the SpectralUNET mapping the initial 238-dimension spectral signatures to 256 dimensions, the spatial-spectral method maps it to 64 dimensions and perhaps causes the model to lose valuable spectral information. Another potential explanation for the limited improvement may be that the band selection method may not be optimal and other metrics should be considered when trying to differentiate root from soil. Finally, the small dataset bounds all three segmentation methods’ performance - a typical issue for deep learning models.

### 6.5. Future Work

The primary path of future work lies in acquiring and publicly releasing additional HSI plant root datasets that detail different species, soil types, soil salinities [56], and soil depths, and by simultaneously gathering other non-destructive imaging datasets (ie. X-ray CTs, MRIs), researchers can improve validation of any HSI-based algorithms and data analytics.

Briefly, additional HSI plant root datasets could allow researchers to take advantage of pretraining methods to improve segmentation performance. It is also possible to use the PRMI dataset for pretraining [12, 57], though some preprocessing or training modifications will likely be needed to make PRMI training like HyperPRI’s RGB data. Another way of improving segmentation performance is with data augmentation. Although well-used for RGB data, it is not always clear what the best approach is for augmenting HSI data. Thus, proper spectral augmentation could help us gain improvements analogous to what occurs when RGB data is properly augmented.

Regarding model-based improvements, larger segmentation architectures (eg. DeepLabv3+ [58]) and recently proposed graph-based methods ([59]) may be explored, but we expect similar conclusions regarding the benefit to segmentation performance afforded by utilizing hyperspectral features in tandem with spatial information. It may also be possible to direct a model’s “attention” toward improving uncertain pixel predictions by adding modules that make the model aware of uncertainty in its encoded features [60]. When used for root segmentation, uncertainty-awareness may decrease noisiness on the root borders or on spectral signatures that are within the overlapping distributions seen in Figure 7. Another proposed path of future work is in the use of texture-based methods to exploit the observation that sections of soil are highly textured, while sections of roots are not as textured and are mostly flat. Conventional convolutional methods are prone to missing small, local statistical changes that are typical of texture. Modifying the models to take advantage of these statistical features can help improve their performance. Finally, we conjecture that because the use of hyperspectral features increased the precision of the deep learning segmentation model, researchers may also expect improvement to proposed root repair algorithms ([61, 62]) and can investigate this utilizing data publicly released through this article.

## 7. Conclusions

We proposed a fully-annotated HSI rhizobox dataset for the peanut and sweet corn annual plants. It provides data that enables researchers to model HS root traits across time and to create learning models meant to target thin, fragmented object features. We detailed the data acquisition methodology, discussed some potential applications, and investigated the use of spatial, spectral, and spatial-spectral methods in a binary segmentation application. Of these three baseline models, the spatial-spectral model demonstrates better overall performance and shows the benefit of using the proposed dataset to add HS features to the spatial model’s input. We briefly described future paths for improving the segmentation performance and recommended possible augmentations for training with the data and with different model variations.

## Supporting information

Graphical Abstract

## Acknowledgements

This research was supported in part by the intramural research program of the U.S. Department of Agriculture, National Institute of Food and Agriculture, Soil Health program under accession no. 1024671. Work conducted by Yangyang Song and William M. Hammond was also supported in part by the Southeastern Peanut Research Initiative and Florida Peanut Producers Association.

## 8. Data and Source Code Availability

As mentioned in the paper, the datasets may be accessed on and downloaded from the following DataVerse link doi: 10.1101/2023.09.29.559614v1.s An associated ‘introductory dataset video’ may be found here: https://youtu.be/T1D1MBxySlI. The source code used for training the UNET, SpectralUNET, and CubeNET models may be found at https://github.com/GatorSense/HyperPRI.

## 9. CRediT Statement

**Spencer J. Chang**: Methodology, Software, Formal Analysis, Investigation, Data Curation, Writing - Original Draft, Visualization. **Ritesh Chowdhry**: Investigation, Data Curation, Writing – Original Draft. **Yangyang Song**: Methodology, Investigation, Data Curation, Writing – Review & Editing. **Tomas Mejia**: Investigation, Data Curation. **Anna Hampton**: Investigation, Data Curation. **Shelby Kucharski**: Writing – Review & Editing. **TM Sazzad**: Writing – Review & Editing. **Yuxuan Zhang**: Writing – Review & Editing. **Sanjeev J. Koppal**: Conceptualization, Writing – Review & Editing. **Chris H. Wilson**: Conceptualization, Writing – Review & Editing. **Stefan Gerber**: Writing – Review & Editing. **Barry Tillman**: Resources, Writing – Review & Editing. **Marcio F. R. Resende**: Resources, Writing – Review & Editing. **William M. Hammond**: Conceptualization, Methodology, Resources, Writing – Review & Editing, Supervision, Project administration. **Alina Zare**: Conceptualization, Methodology, Resources, Writing – Review & Editing, Supervision, Project administration, Funding acquisition.

## Appendix A. Additional HSI Phenotype Comparisons and Data Splits

**Table A.6:** Split counts for each day in each peanut dataset split. Legend is (#train/#val).

| Date | Split 1 | Split 2 | Split 3 | Split 4 | Split 5 |
| --- | --- | --- | --- | --- | --- |
| **Jun-24** | 13/5 | 16/2 | 10/8 | 16/2 | 14/4 |
| **Jul-1** | 8/3 | 10/1 | 6/5 | 9/2 | 9/2 |
| **Jul-8** | 3/1 | 3/1 | 4/0 | 2/2 | 2/2 |
| **Jul-14** | 4/1 | 4/1 | 4/1 | 3/2 | 3/2 |
| **Jul-21** | 2/1 | 2/1 | 3/0 | 2/1 | 1/2 |
| **Jul-27** | 3/1 | 3/1 | 4/0 | 2/2 | 2/2 |
| **Aug-6** | 3/1 | 3/1 | 4/0 | 2/2 | 2/2 |
| **Aug-8** | 2/0 | 1/1 | 2/0 | 1/1 | 2/0 |
| **Aug-10** | 2/0 | 1/1 | 2/0 | 1/1 | 2/0 |
| **Aug-12** | 1/0 | 0/1 | 1/0 | 1/0 | 1/0 |
| **Aug-15** | 1/0 | 0/1 | 1/0 | 1/0 | 1/0 |
| **Aug-19** | 1/0 | 0/1 | 1/0 | 1/0 | 1/0 |
| **Aug-21** | 1/0 | 0/1 | 1/0 | 1/0 | 1/0 |
| **Aug-24** | 1/0 | 0/1 | 1/0 | 1/0 | 1/0 |
| **Total** | **45/14** | **44/15** | **45/14** | **44/15** | **43/16** |

**Figure A.15:**
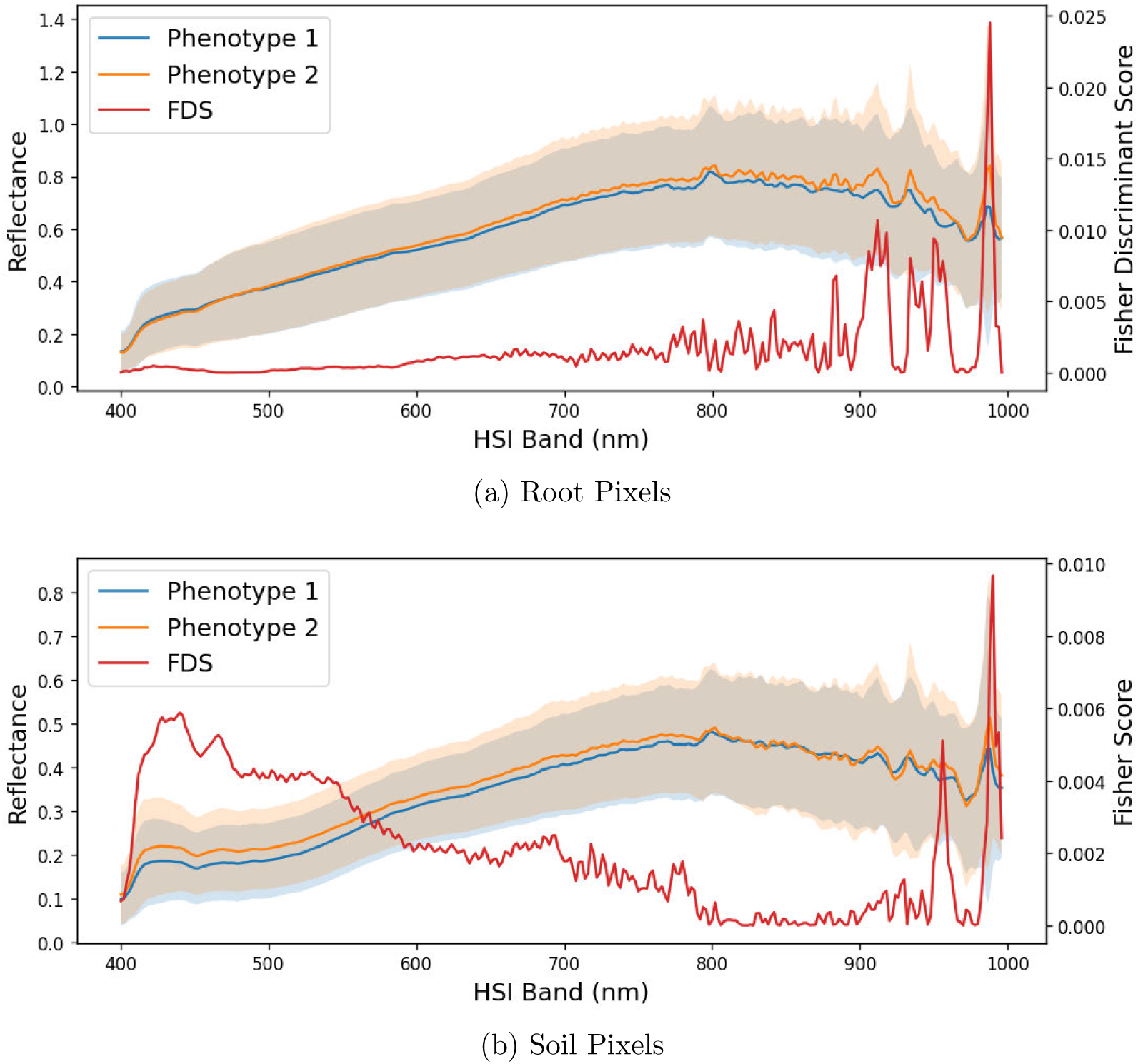
Additional comparison of data spectra and edges for peanut phenotypes 1 and 2. (a) and (b) show the FDS between soil and root, respectively, are around 10x less than the FDS of soil versus root pixels.

## Appendix B. Selected Segmentation Results

In the following figures, selected segmentation results are chosen for com parison between UNET, SpectralUNET, and CubeNET. As of the writing and analysis presented in this paper, the image for 20220806 box53 was mislabeled, so all models missed the same peanut peg in their own ways. As mentioned before in Fig. 11, darkened image pixels are true negatives. Red pixels are false positives. Blue pixels are false negatives. Yellow-green pixels are true positives.

**Figure B.16:**
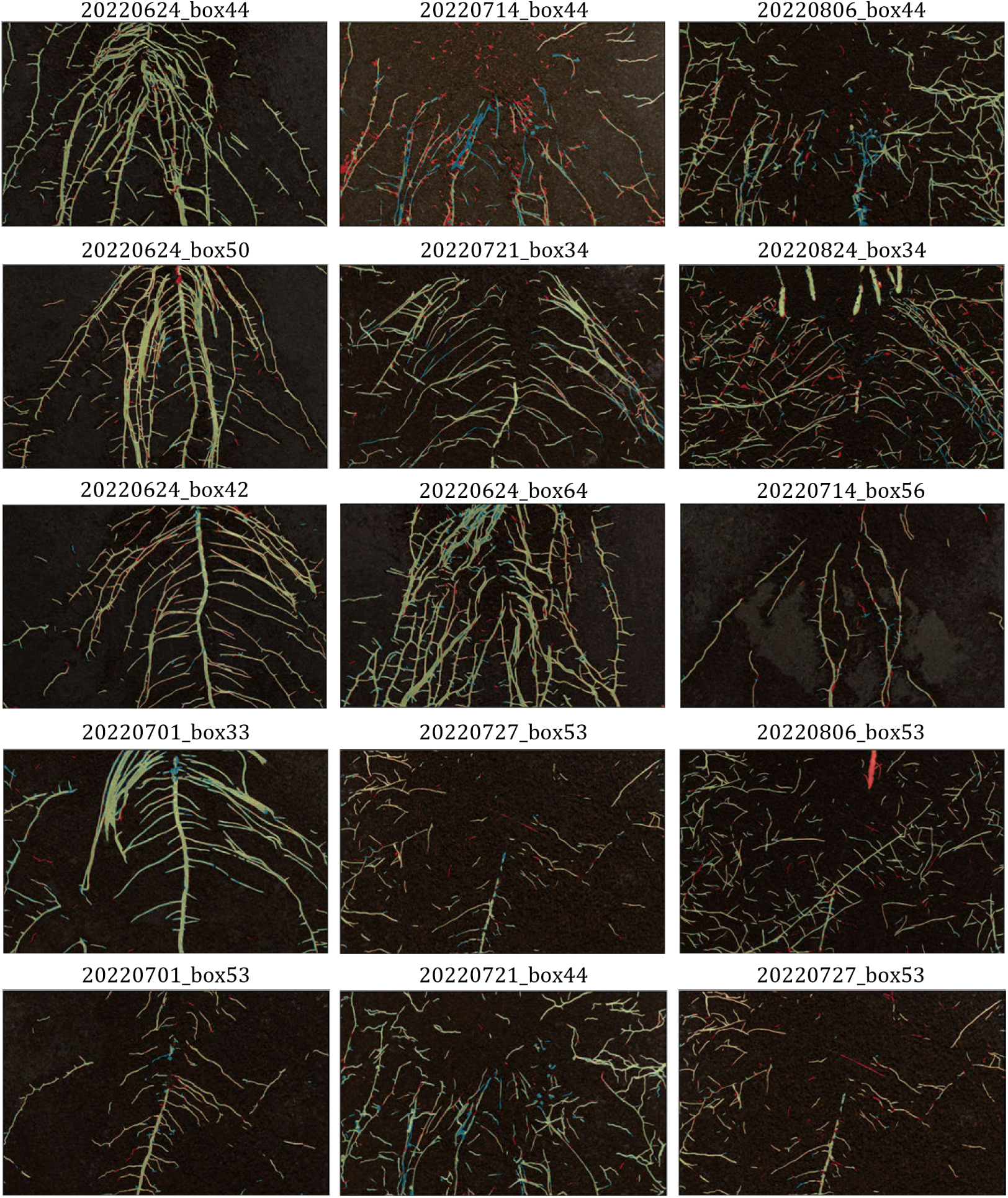
Selected segmentation results for UNET. Each row corresponds to one split of data (ie. the second row takes segmentations from split 2).

**Figure B.17:**
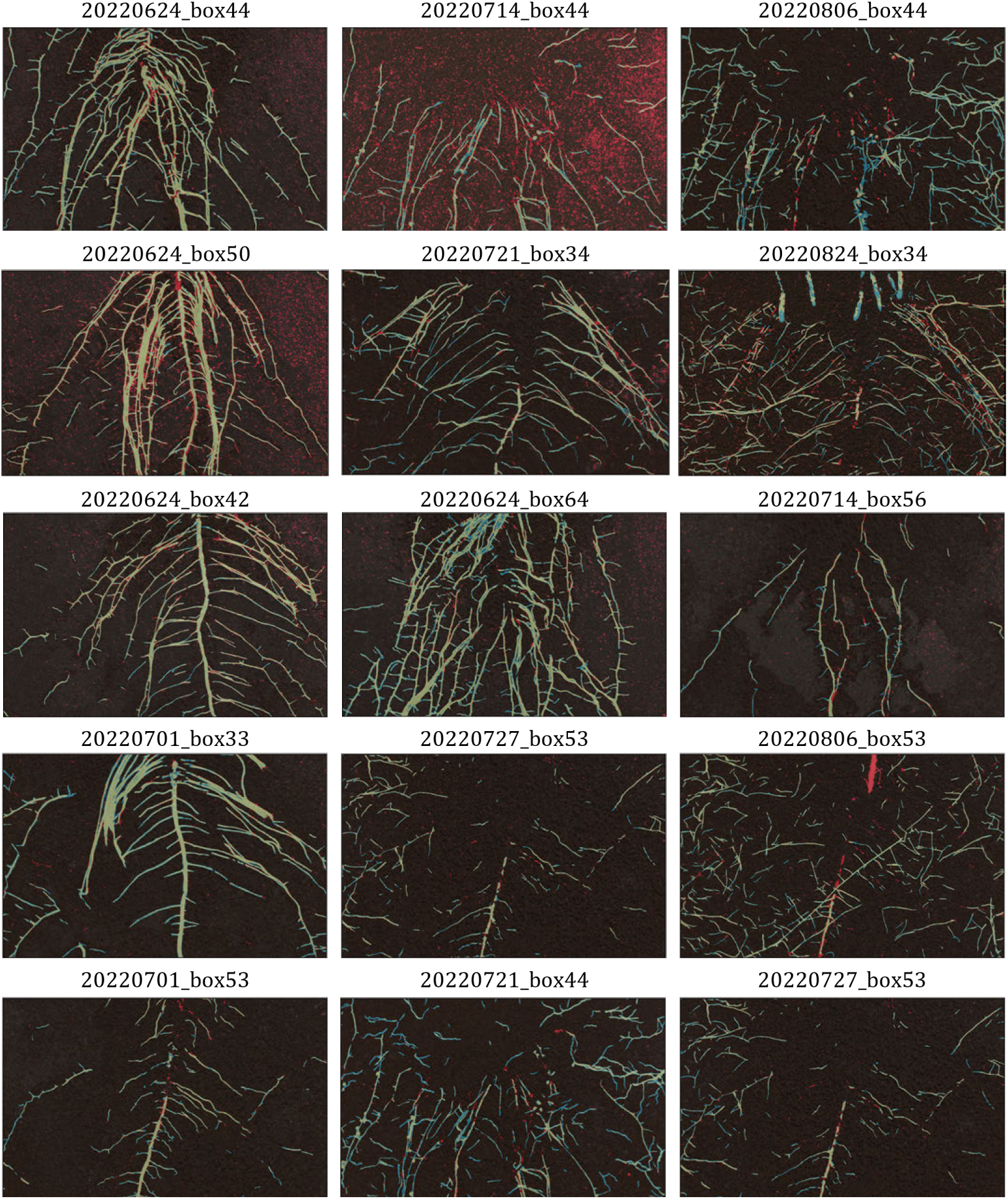
Selected segmentation results for SpectralUNET. Each row corresponds to one split of data (ie. the second row takes segmentations from split 2).

**Figure B.18:**
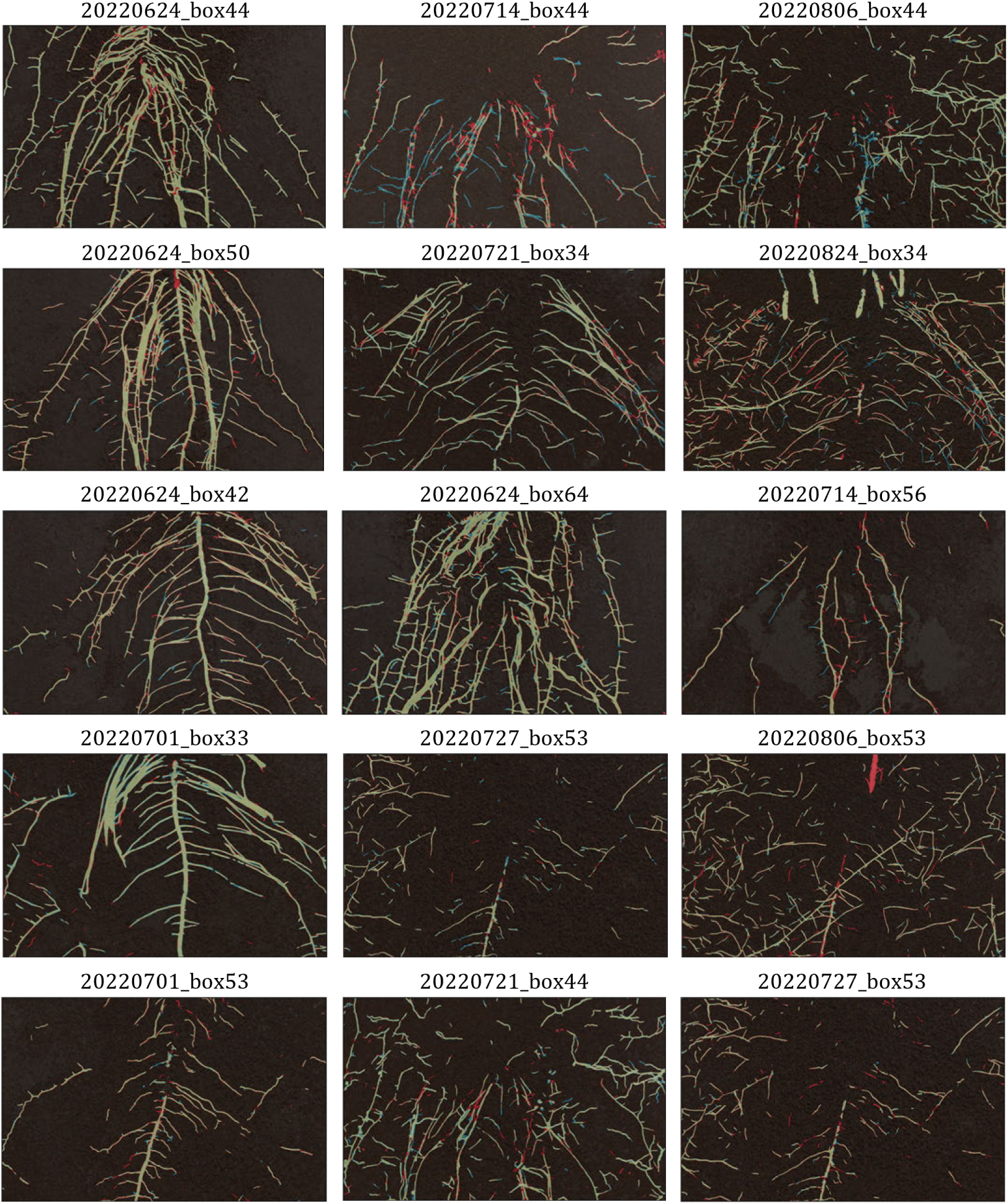
Selected segmentation results for CubeNET. Each row corresponds to one split of data (ie. the second row takes segmentations from split 2).

