## Supplementary material for "HyperPRI: A Dataset of Hyperspectral Images for Underground Plant Root Study": Graphical Abstract

Enhance **plant root analyses** and tackle **challenging features with machine learning** models.

https://linktr.ee/spjchang

**Without Hyperspectral Information** **With Hyperspectral Information**

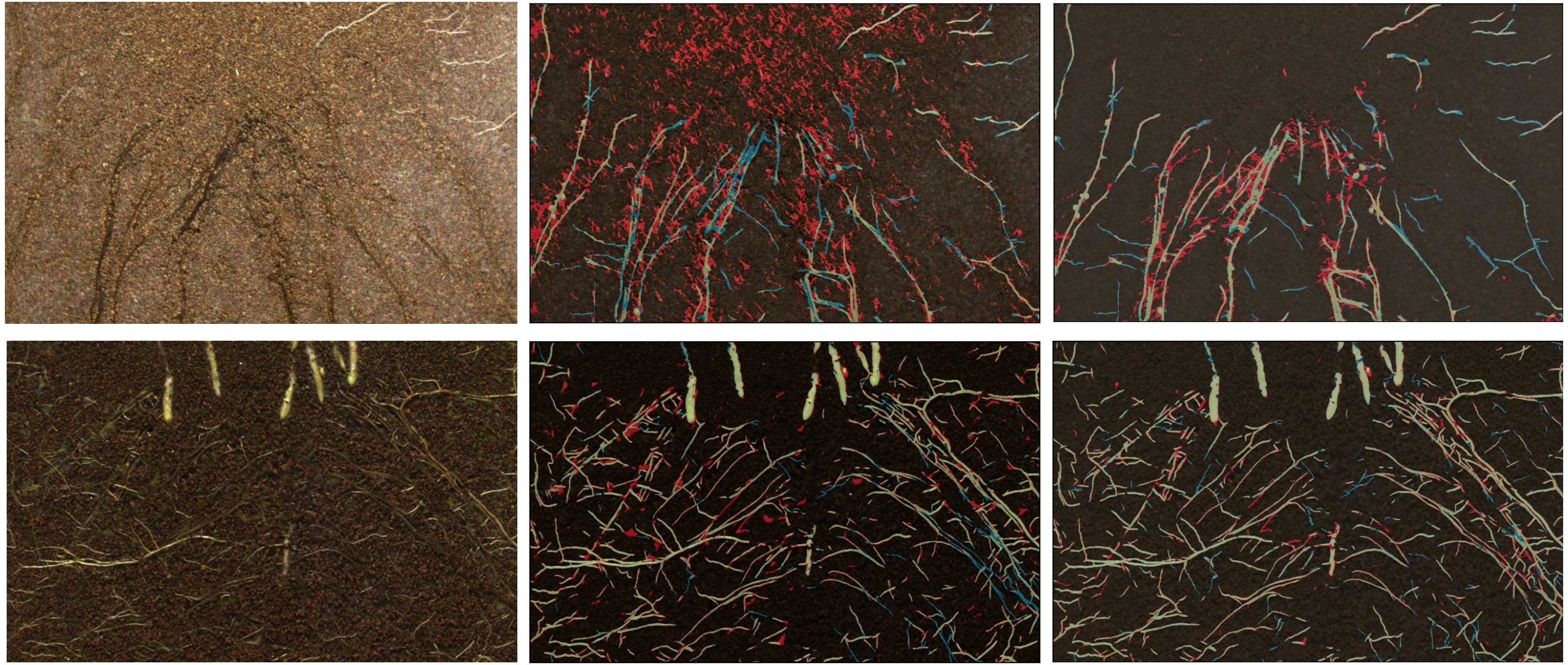

Color Legend: Correctly Classified Root | Misclassified Soil | Misclassified Root

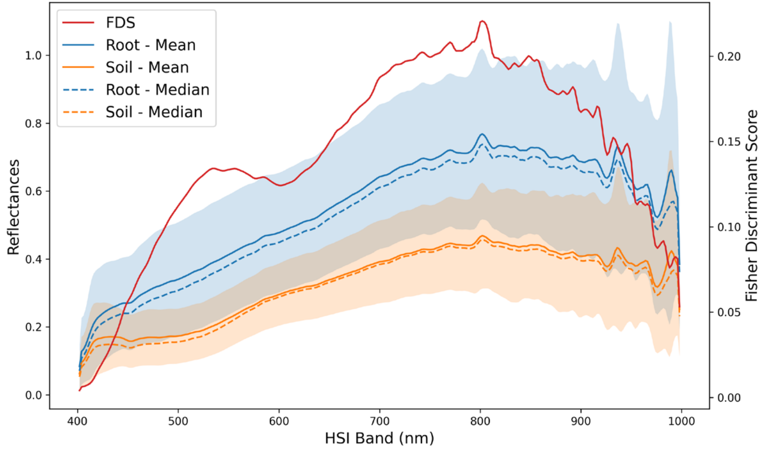


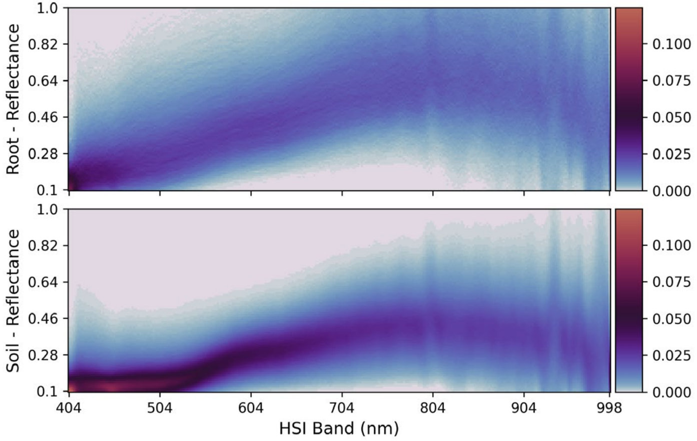



Spencer J Chang, Ritesh Chowdhry, Yangyang Song, et al (2023)

THE MACHINE LEARNING AND SENSING LABORATORY
ecophyslab
FOCUS
DEPARTMENT OF ELECTRICAL AND COMPUTER ENGINEERING
